# Rapid protection from COVID-19 in nonhuman primates vaccinated intramuscularly but not intranasally with a single dose of a recombinant vaccine

**DOI:** 10.1101/2021.01.19.426885

**Authors:** Wakako Furuyama, Kyle Shifflett, Amanda N. Pinski, Amanda J. Griffin, Friederike Feldmann, Atsushi Okumura, Tylisha Gourdine, Allen Jankeel, Jamie Lovaglio, Patrick W. Hanley, Tina Thomas, Chad S. Clancy, Ilhem Messaoudi, Kyle L. O’Donnell, Andrea Marzi

## Abstract

The ongoing pandemic of Coronavirus disease 2019 (COVID-19) continues to exert a significant burden on health care systems worldwide. With limited treatments available, vaccination remains an effective strategy to counter transmission of severe acute respiratory syndrome coronavirus 2 (SARS-CoV-2). Recent discussions concerning vaccination strategies have focused on identifying vaccine platforms, number of doses, route of administration, and time to reach peak immunity against SARS-CoV-2. Here, we generated a single dose, fast-acting vesicular stomatitis virus-based vaccine derived from the licensed Ebola virus (EBOV) vaccine rVSV-ZEBOV, expressing the SARS-CoV-2 spike protein and the EBOV glycoprotein (VSV-SARS2-EBOV). Rhesus macaques vaccinated intramuscularly (IM) with a single dose of VSV-SARS2-EBOV were protected within 10 days and did not show signs of COVID-19 pneumonia. In contrast, intranasal (IN) vaccination resulted in limited immunogenicity and enhanced COVID-19 pneumonia compared to control animals. While IM and IN vaccination both induced neutralizing antibody titers, only IM vaccination resulted in a significant cellular immune response. RNA sequencing data bolstered these results by revealing robust activation of the innate and adaptive immune transcriptional signatures in the lungs of IM-vaccinated animals only. Overall, the data demonstrates that VSV-SARS2-EBOV is a potent single-dose COVID-19 vaccine candidate that offers rapid protection based on the protective efficacy observed in our study.

**One sentence summary:** VSV vaccine protects NHPs from COVID-19 in 10 days

## Introduction

Severe acute respiratory syndrome coronavirus 2 (SARS-CoV-2) is a positive-sense, single-stranded RNA virus first isolated from a patient with severe respiratory illness in Wuhan, China (*1*). SARS-CoV-2 infection manifests as a clinical syndrome termed Coronavirus disease 2019 (COVID-19), which can lead to respiratory failure (*2*). In addition to respiratory distress, other clinical manifestations associated with SARS-CoV-2 infection include cardiac pathology, gastrointestinal disease, coagulopathy, and hyperinflammatory syndrome (*3-5*). Patients with an increased risk of severe clinical manifestation include the elderly, immunocompromised, and individuals with co-morbidities (obesity, diabetes, hypertension etc.)(*6*). Virtually every country has been affected with almost one hundred million infections to date and an estimated case fatality rate of ∼2% (https://coronavirus.jhu.edu/map.html). The widespread morbidity, mortality, and socioeconomic impact of COVID-19 emphasize the urgent need for the development and deployment of countermeasures, including vaccines.

The COVID-19 pandemic has made the development of a vaccine a global priority (*7-9*). An ideal vaccine candidate is safe, effective, fast-acting, rapidly deployable, and requires only a single immunization. Most of the current vaccine candidates encode the trimeric SARS-CoV-2 spike (S) protein as the primary antigen. S is essential for SARS-CoV-2 infectivity since it binds the angiotensin-converting enzyme 2 (ACE2) receptor and promotes viral-cell membrane fusion (*10*). It is also the main target for virus neutralization (*11*). The route of vaccination can greatly influence the local immune environment at the vaccination and infection site. Recently, the comparison of intramuscular (IM) and intranasal (IN) vaccination of mice with a chimpanzee adenoviral vector-based vaccine revealed an increase in stimulation of local mucosal immunity and generation of antigen-specific IgA and lung resident T cells after IN vaccination. The local mucosal immunity was improved by the generation of antigen-specific IgA and lung resident T cell generation after IN vaccination (*12*). Prior to progression to human clinical trials, several COVID-19 vaccine candidates were IM administered to nonhuman primates (NHPs) to evaluate their efficacy (*13-16*).

The recombinant vesicular stomatitis virus (VSV) vaccine platform has previously been utilized in vaccines against multiple viral pathogens, such as Ebola virus (EBOV), Marburg, Nipah, and Lassa viruses (*17-19*). VSV-based vaccines elicit a robust and rapid immune response to the encoded antigen(s) after a single immunization (*20*). The time to immunity has been demonstrated to be within 7-10 days for a number of pathogens in preclinical and clinical studies, greatly reducing the time needed between vaccination and protection (*21-24*). Multiple routes of VSV-based vaccination, such as IM and IN, have been shown to be efficacious (*21, 22, 25*). Furthermore, the general population is predominantly seronegative for VSV, circumventing pre-exisiting immunity neutralizing the vaccine virus (*20*). These unique attributes - robust immune stimulation and time to immunity - make this an attractive vaccine platform for SARS-CoV-2. However, the immunogenicity and efficacy of an IM- or IN-administered COVID-19 VSV-based vaccine has not been tested in the NHP model.

In the present study, we developed a VSV-based vector expressing the SARS-CoV-2 S in combination with the EBOV glycoprotein (GP). We utilized the NHP challenge model and compared the vaccine efficacy with a shorter time to challenge in tandem with comparing the optimal route of immunization. We demonstrate that IM-vaccinated NHPs developed no to mild lesions of COVID-19 with variable immunopathology, whereas IN vaccination resulted immune-enhanced disease with interstitial pneumonia in NHPs. IM vaccination resulted in robust and rapid humoral and cellular immune responses while with IN vaccination did not. Transcriptional analysis of the lungs supports our immunological findings by revealing greater expression of innate and adaptive immune genes in the IM vaccination group.

## Results

### Vaccine construction and characterization

The VSV-backbone encoding the EBOV Kikwit GP, rVSV-ZEBOV, was used as a parental vector to construct this COVID-19 vaccine. Therefore, we generated a VSV construct co-expressing the EBOV GP and SARS-CoV-2 S (VSV-SARS2-EBOV) by the adding the full-length codon-optimized SARS-CoV-2 S upstream of the EBOV GP into the existing VSV vector (Fig. S1A). The construct was recovered from plasmid following previously established protocols (*26*). Expression of both antigens, SARS-CoV-2 S and EBOV GP, was confirmed by Western blot analysis of the VSV particles in cell supernatant (Fig. S1B). Next, we performed viral growth kinetics. VSV-SARS2-EBOV replicated with similar kinetics and had comparable endpoint titers to the parental VSV-EBOV in Vero E6 cells (Fig. S1C), which does not impact potential vaccine production.

### Efficacy in nonhuman primates (NHPs)

We demonstrated previously that the parental VSV-EBOV is a fast-acting, IM-administered vaccine (*21*); that was confirmed in human phase 3 clinical trials (*24*). Therefore, we set out to analyze the fast-acting potential of a single dose of this VSV-based COVID-19 vaccine in rhesus macaques. Unlike EBOV disease (EVD), COVID-19 is a respiratory disease. However, a previously published study demonstrated that mucosal immunization with VSV-EBOV protected the NHPs from EVD (*25*). To determine the efficacy of mucosal vaccination against COVID-19 in NHPs, we compared the efficacy of IM and IN vaccination in the rhesus macaque model (*27*). Groups of 6 NHPs were either IN- or IM-vaccinated with VSV-SARS2-EBOV while control animals received a single dose of VSV-EBOV IN (n=2) or IM (n=2)(Fig. S1D). All NHPs were observed for potential adverse effects, particularly after IN vaccination as this is not the standard route of administration for this vaccine platform, but no clinical changes were noted. After 10 days, all NHPs were challenged with SARS-CoV-2 as previously described (*27*). On days post challenge (dpc) 0, 1, 3, 5, and 7, a clinical exam including thoracic radiographs and nasal swab collection was performed; in addition, the dpc 3 exam included a bronchoalveolar lavage (BAL). On dpc 7, all NHPs were euthanized and samples were collected for analysis. None of the vaccinated animals displayed clinical signs of disease after challenge.

We determined differences in total SARS-CoV-2 RNA and subgenomic (sg) RNA in the nasal swabs of the animals throughout the study. Interestingly, IN vaccination resulted in significantly lower levels of nasal viral RNA on dpc 1 compared to IM indicating better local control of virus replication (Fig. 1A). However, on dpc 3 only the total SARS-CoV-2-specific RNA levels were significantly different (Fig.1A). In contrast, both total and sg RNA levels in the BAL were significantly lower for the IM vaccination group compared to IN-vaccinated and control groups (Fig. 1B). This finding is supported by our observation that the IN-vaccinated NHPs had more lung infiltrates compared to the IM and control groups at the time of euthanasia (Fig. 1C). Additionally, only IN-vaccinated NHPs exhibited lung lesions (Fig. S2). This was accompanied by a significant reduction of total RNA and sgRNA in IM-vaccinated, but not IN-vaccinated NHPs compared to controls (Fig. 1D). The comparison of RNA levels in individual lung lobes and other examined tissue samples did not reveal any significant differences (Fig. S3).

**Figure 1.**
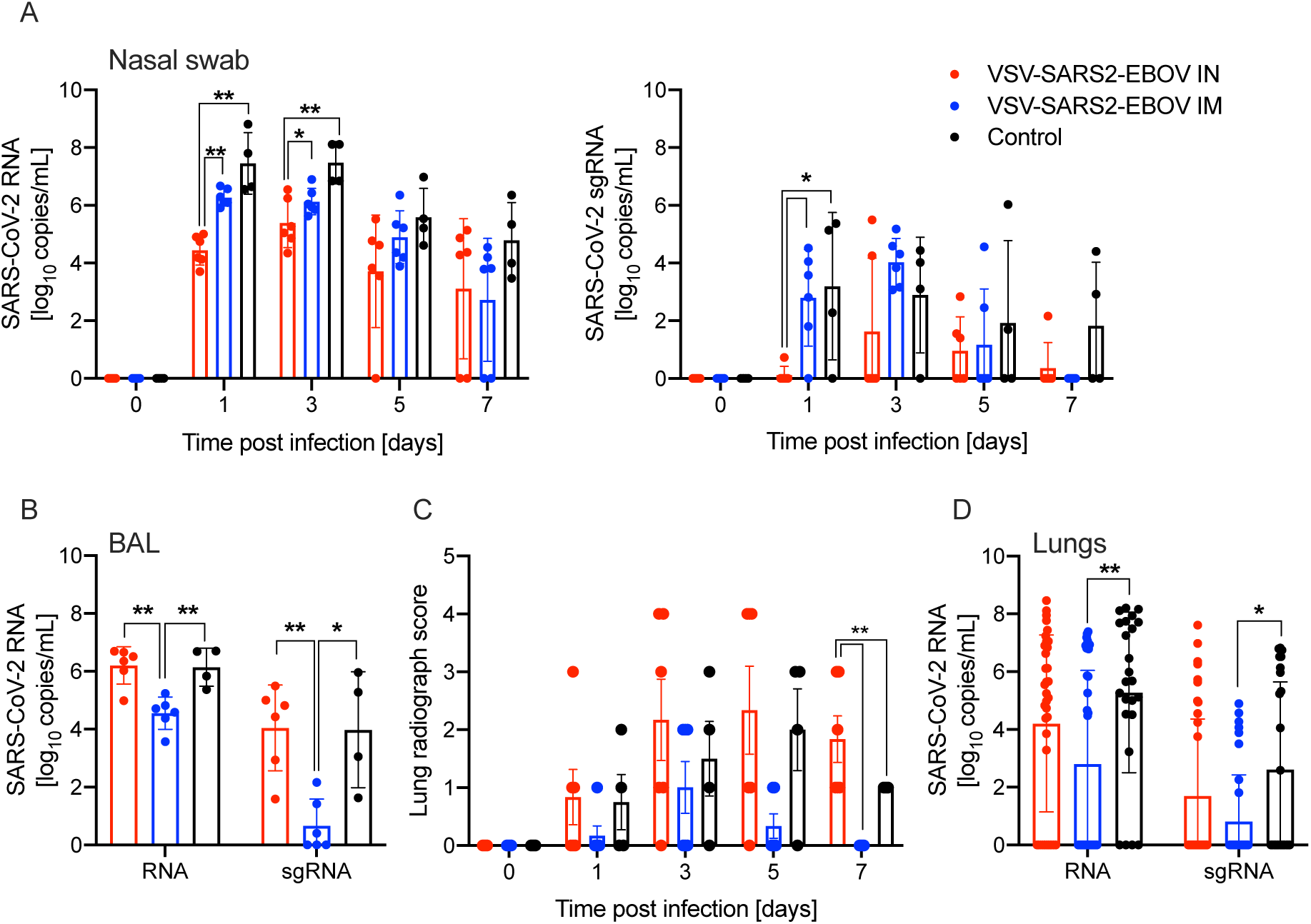
SARS-CoV-2 loads in vaccinated NHPs. Groups of 6 NHPs were IN or IM vaccinated with a single dose of VSV-SARS2-EBOV; 4 control animals received the VSV-EBOV. **(A)** Total SARS-CoV-2-specific RNA (left panel) and subgenomic (sg) RNA (right panel) in nasal swabs collected from NHPs. **(B)** Total SARS-CoV-2-specific RNA and sgRNA in bronchoalveolar lavage (BAL) samples collected on day 3. **(C)** Lung radiograph scores after challenge. Mean and standard deviation (SD) are shown. **(D)** Total SARS-CoV-2-specific RNA and sgRNA in lung samples collected on day 7. **(A, B, D)** Geometric mean and geometric SD are depicted. Statistical significance is indicated.

### Histopathology in NHPs

Histopathologic analysis of the collected lung samples revealed pulmonary pathology consistent with the previously described rhesus macaque model of SARS-CoV-2 infection in the control group regardless of IM or IN administration of the control vaccine (Fig. 2A,D)(*27*). In the IM vaccination group pulmonary lesions consisted of low to moderate numbers of eosinophils multifocally infiltrating bronchiolar mucosa, excess mucus accumulation in the lumen of bronchi and bronchioles, and profound perivascular lymphocytic cuffing (interpreted as immune pathology) disseminated throughout all lung lobes (Fig. 2B,E). In combination, these lesions are suggestive of a localized hypersensitivity response.

**Figure 2.**
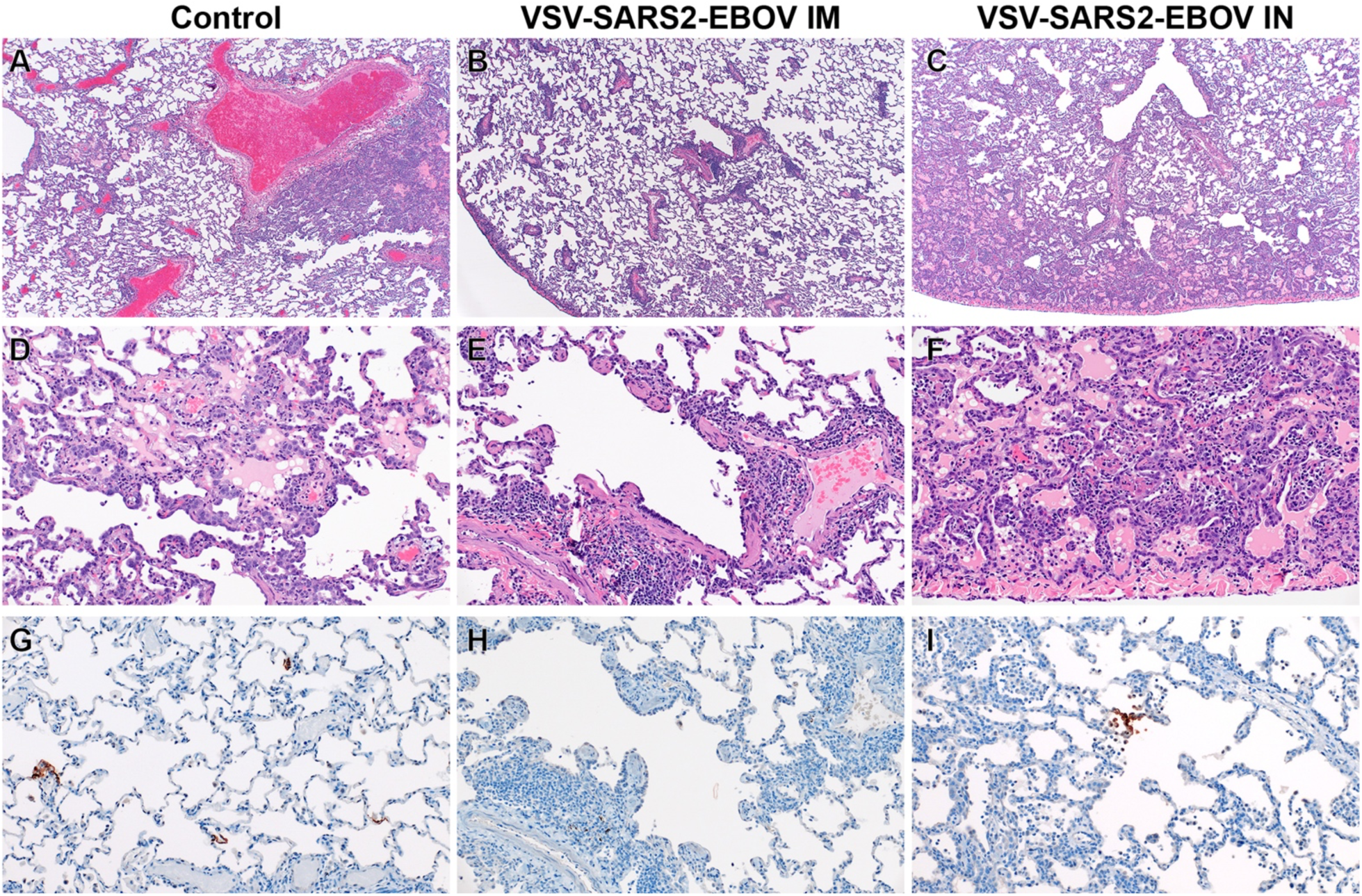
Histopathology and Immunohistochemistry of NHP lungs. **(A)** Pulmonary lesions depicting typical coronavirus respiratory pathology including locally extensive regions of bronchointerstitial pneumonia and proteinaceous fluid accumulation in adjacent alveoli (40x, H&E). **(B)** Disseminated immunopathology with prominent perivascular lymphocytic cuffing and multifocal involvement at terminal airways (40x, H&E). **(C)** IN vaccination shows pulmonary pathology characterized by a combination of interstitial pneumonia and immunopathology (40x, H&E). **(D)** Foci of interstitial pneumonia are characterized by prominent type II pneumocyte hyperplasia, leukocyte infiltration and expansion of alveolar septa and accumulation of low numbers of macrophages, neutrophils and proteinaceous fluid in alveolar spaces (200x, H&E). **(E)** Terminal airways and medium to small caliber blood vessels are cuffed by moderate numbers of lymphocytes with scattered eosinophils (200x, H&E). **(F)** Foci of interstitial pneumonia show pronounced type II pneumocyte hyperplasia, thickening of alveolar septa by an infiltration of leukocytes and leukocyte spillover into adjacent alveolar spaces with moderate numbers of alveolar eosinophils noted and multifocal fibrin mats filling alveolar spaces (200x, H&E). **(G)** Low numbers of type I pneumocytes in regions lacking pathology are immunoreactive for SARS-CoV-2 antibody (200x, immunohistochemistry (IHC)). **(H)** SARS-CoV-2-specific immunoreactivity was not observed in evaluated sections of the IM vaccinated group (200x, IHC). **(I)** Low numbers of type I pneumocytes and alveolar macrophages are immunoreactive for SARS-CoV-2 in select foci of interstitial pneumonia (200x, IHC).

Hypersensitivity lesion location mirrored that of what has previously been described in this NHP model of COVID-19 (*27*), with a high proportion of lesions located at the periphery of the lung and increased lesion severity in lower lung lobes. Limited evidence of type I pneumocyte damage was present in rare foci and was characterized by lining of alveoli by type II pneumocytes and a scant amount of proteinaceous fluid within alveolar spaces (Fig. 2B,E). Histopathologic lesions in the IN vaccination group mirrored the enhanced gross lesion severity and histologically consisted of an immune-enhanced disease with evidence of classic moderate to severe SARS-CoV-2 pulmonary pathology and moderate hypersensitivity response (Fig. 2C,F). The hypersensitivity response was similar to that observed in the IM-vaccinated group, but more severe with the addition of eosinophil spillover into bronchiolar lumen and moderate numbers of alveolar spaces. SARS-CoV-2 nucleoprotein immunoreactivity was observed in type I pneumocytes and macrophages of both the control and IN groups, but not in the IM vaccination group (Fig. 2G-I).

### Immune responses in NHPs

We next analyzed the peripheral humoral response. IgG responses to the full-length SARS-CoV-2 S, the S receptor binding domain (RBD), and the EBOV GP were determined following vaccination and challenge (Fig. 3). We demonstrated that the IM-vaccinated NHPs attained significantly higher SARS-CoV-2 S-specific IgG starting 10 days after vaccination and following challenge compared to IN-vaccinated NHPs and controls (Fig. 3A). Similarly, the IgG response to the SARS-CoV-2 S RBD was higher at day 10 post vaccination (0 dpc) in the IM group compared to the IN and controls (Fig. 3B). Analysis of the IgG subclasses in serum on dpc 0 and 7 showed that both vaccination routes resulted in predominantly IgG1, IgG2 and IgG3 antibodies with no significant difference between the groups (Fig. 3C). Only IgG1 increased after challenge in both vaccine groups significantly compared to controls (Fig. 3C). Interestingly, on dpc 7 both IgG2 and IgG3 antibodies were largely absent in sera from IN-vaccinated NHPs (Fig. 3C). IgG responses specific to EBOV GP support the finding that IM vaccination appears more immunogenic compared to IN even though the data are only significantly different on day 0 and day 3 (Fig. S4A). Measurable SARS-CoV-2 neutralizing titers were detected as early as 10 days following vaccination for the IM group, and 11 days for the IN group (Fig. 3D). On day 3, significantly higher titers were observed comparing IM-vaccinated NHPs to controls only (Fig. 3D). At the time of euthanasia, the neutralizing titers in IM- and IN-vaccinated NHPs were comparable but significantly higher than those observed in control animals (Fig. 3D)

**Figure 3.**
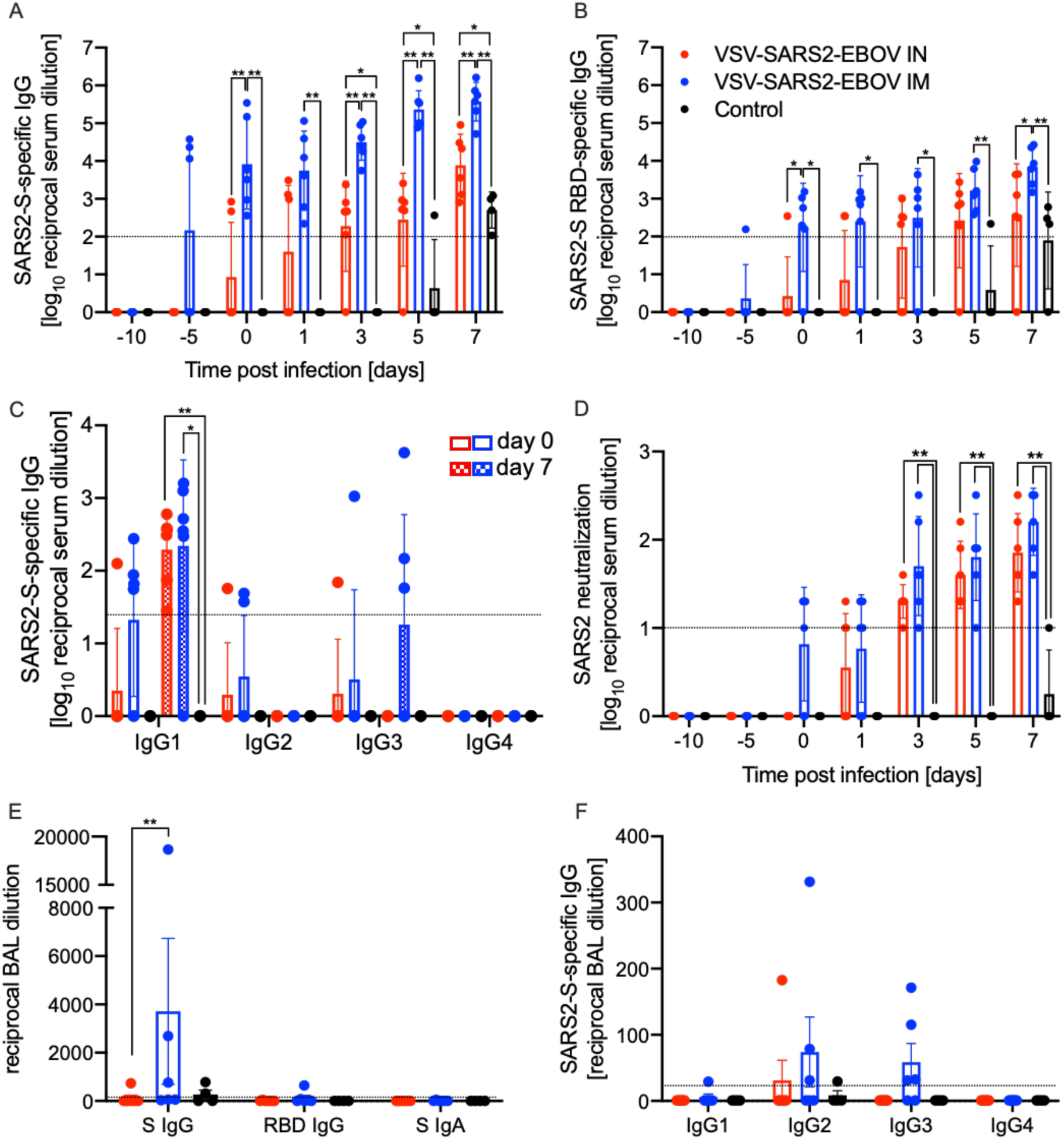
Humoral immune responses in NHPs. Serum samples collected throughout the study from all NHPs were examined for **(A)** SARS-CoV-2 S-specific IgG, **(B)** SARS-CoV-2 S receptor binding domain (RBD)-specific IgG or **(C)** IgG subclasses specific to SARS-CoV-2 S by ELISA. **(D)** Neutralizing titers to SARS-CoV-2 were determined. **(E)** Bronchoalveolar lavage (BAL) samples were analyzed for SARS-CoV-2 S-specific IgG (S IgG) or IgA (S IgA), and SARS CoV-2 S RBD-specific IgG (RBD IgG) by ELISA. **(A-D)** Geometric mean and geometric standard deviation (SD) are depicted. **(F)** IgG subclasses specific to SARS-CoV-2 S in BAL samples were analyzed by ELISA. **(E, F)** Mean and SD are depicted. Statistical significance is indicated.

Next, we investigated the humoral responses in the BAL obtained on dpc 3. We detected SARS-CoV-2 S-specific IgG in 3 of the 6 NHPs in the IM vaccinated group but only in 1 of 6 NHPs in the IN group (Fig. 3E). Only the IM-vaccinated NHP with the highest titer of SARS-CoV-2 S-specific IgG had anti-SARS-CoV-2 S RBD IgG (Fig. 3E). SARS-CoV-2 S-specific IgA was not detected in any of the BAL samples (Fig. 3E). In contrast to serum, no IgG1 was detected; however, IgG2 and IgG3 were readily detected in several IM-vaccinated NHPs (Fig. 3F). Unexpectedly, IM vaccination resulted in higher humoral responses in the lung to all antigens including EBOV GP (Fig. S4B).

We analyzed the peripheral cellular response even though T cell responses have shown to play only a limited role mediating protection using the VSV-EBOV vaccine (*28, 29*). Peripheral blood mononuclear cells (PBMCs) were stimulated with a peptide-pool spanning the entire length of the SARS-CoV-2 S and the antigen-specific T cells were identified using intracellular cytokine staining (Fig.4A,B). While there was minimal cytokine production by CD4^+^ T cells, a significant increase in the CD69 activation marker was seen in IM-vaccinated NHPs on 0 and 7 dpc relative to controls (Fig. 4A). Similarly, a significantly higher portion of the CD8^+^ T cells from the IM group produced granzyme B on dpc 0 and 7 compared to controls and IN groups (Fig. 4B). Interestingly, numbers of IL-2^+^ CD4^+^ and CD8^+^ T cells were significantly lower in the IM group on dpc 0, and comparable to IN and control groups by dpc 7 (Fig. 4A,B). Additionally, a greater number of granzyme B+ NK cells was measured on dpc 0 and 7 in IM-vaccinated animals compared to IN-vaccinated and control animals (Fig. 4C).

**Figure 4.**
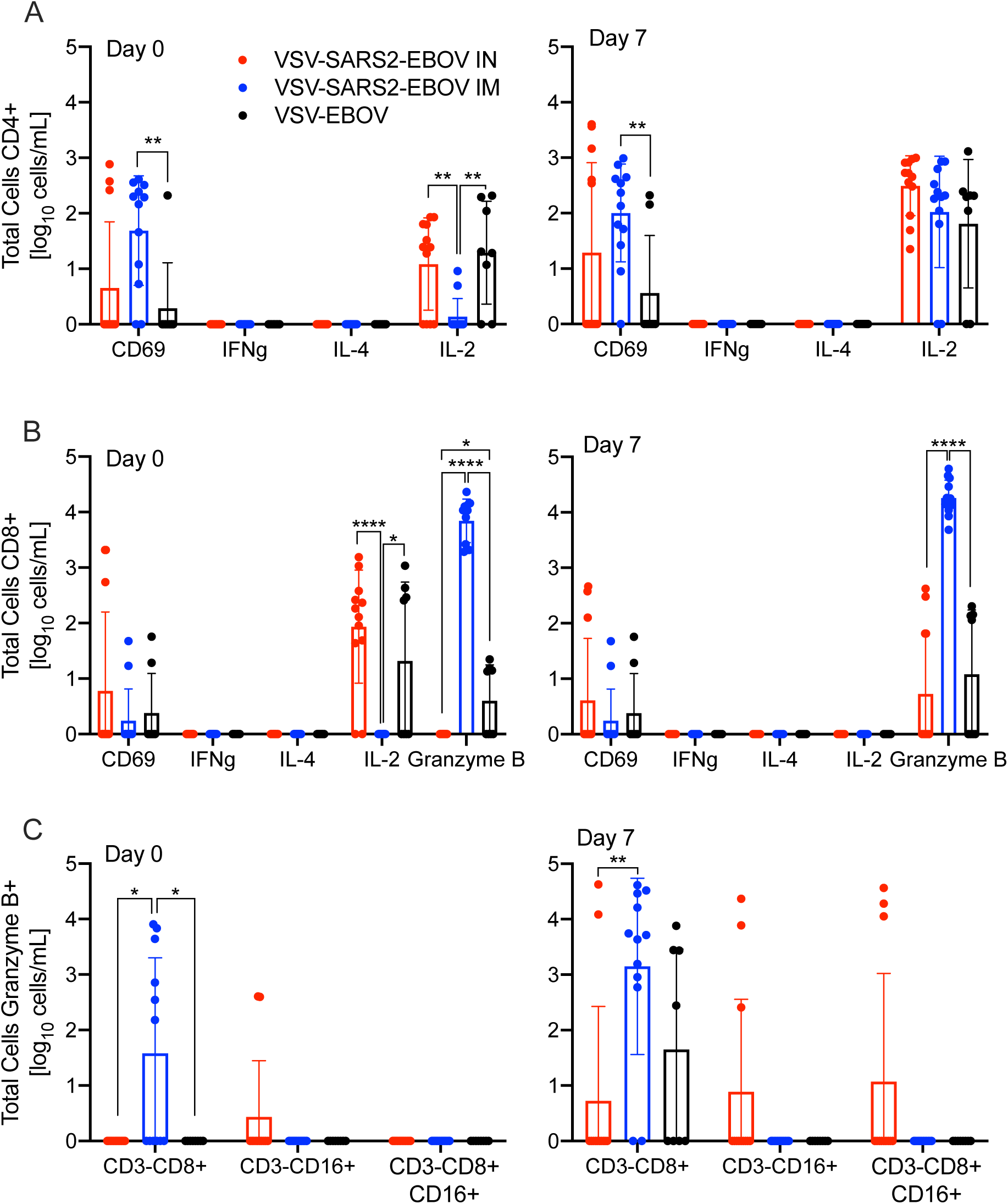
Peripheral cellular immune response post challenge. **(A)** CD4^+^ T cells and from PBMCs were stained for expression of early activation marker CD69 and intracellular cytokine staining (ICS) for IFN γ, IL-4, and IL-2 on day 0 and 7 post challenge. **(B)** CD8^+^ T cells from PBMCs were phenotyped for expression of early activation marker CD69 and ICS for IFN γ, IL-4, IL-2, and granzyme B on day 0 and 7 post challenge. **(C)** NK cell subpopulations were stained for the expression of granzyme B on day 0 and 7 post challenge. Data was measured in duplicate for all animals. Geometric mean and SD are depicted. Statistical significance is indicated.

Finally, we monitored levels of systemic and BAL cytokines and chemokines. We found a significant increase in MCP-1 on dpc 1 and 3 in IN-vaccinated animals compared to IM-vaccinated animals. A similar trend was observed for IL-18 on dpc 3 and 5 (Fig. S5A). Levels of MIP-1β were significantly lower at all dpcs in IM-vaccinated animals compared to the control and IN-vaccinated animals (Fig. S5A). Analysis of cytokine and chemokine levels in the BAL revealed that MCP-1 levels were significantly decreased in IN-vaccinated NHPs (Fig. S5B). All other investigated cytokines did not show significant differences.

### Transcriptional analysis of BAL samples

To better understand the molecular underpinnings of differential vaccine responses, we profiled the host transcriptional response in the BAL (dpc 3) samples. Principal component analysis (PCA) of BAL samples indicated a distinct separation of uninfected/naЇve (historical data) and the three challenged groups, with no clear distinction between the challenged groups (Fig. S6A). Therefore, the samples from control, IM- or IN-vaccinated animals were compared to the uninfected samples. Over 1,000 differentially expressed genes (DEGs) were detected in the challenged groups compared to the uninfected animals, with most DEGs being upregulated (Fig. S6B). The majority of DEGs were shared amongst the three challenged groups (Fig. 5A). Functional enrichment showed that upregulated and downregulated DEGs shared by all three challenged groups play a role in regulating cell structure (e.g., “actin cytoskeleton organization”) and innate immunity (e.g., “positive regulation of cytokine production”, “myeloid leukocyte activation”) (Fig. 5B). However, only upregulated DEGs enriched to gene ontology (GO) terms associated with adaptive immunity (e.g., “lymphocyte activation”, “T cell differentiation”)(Fig. 5B).

**Figure 5.**
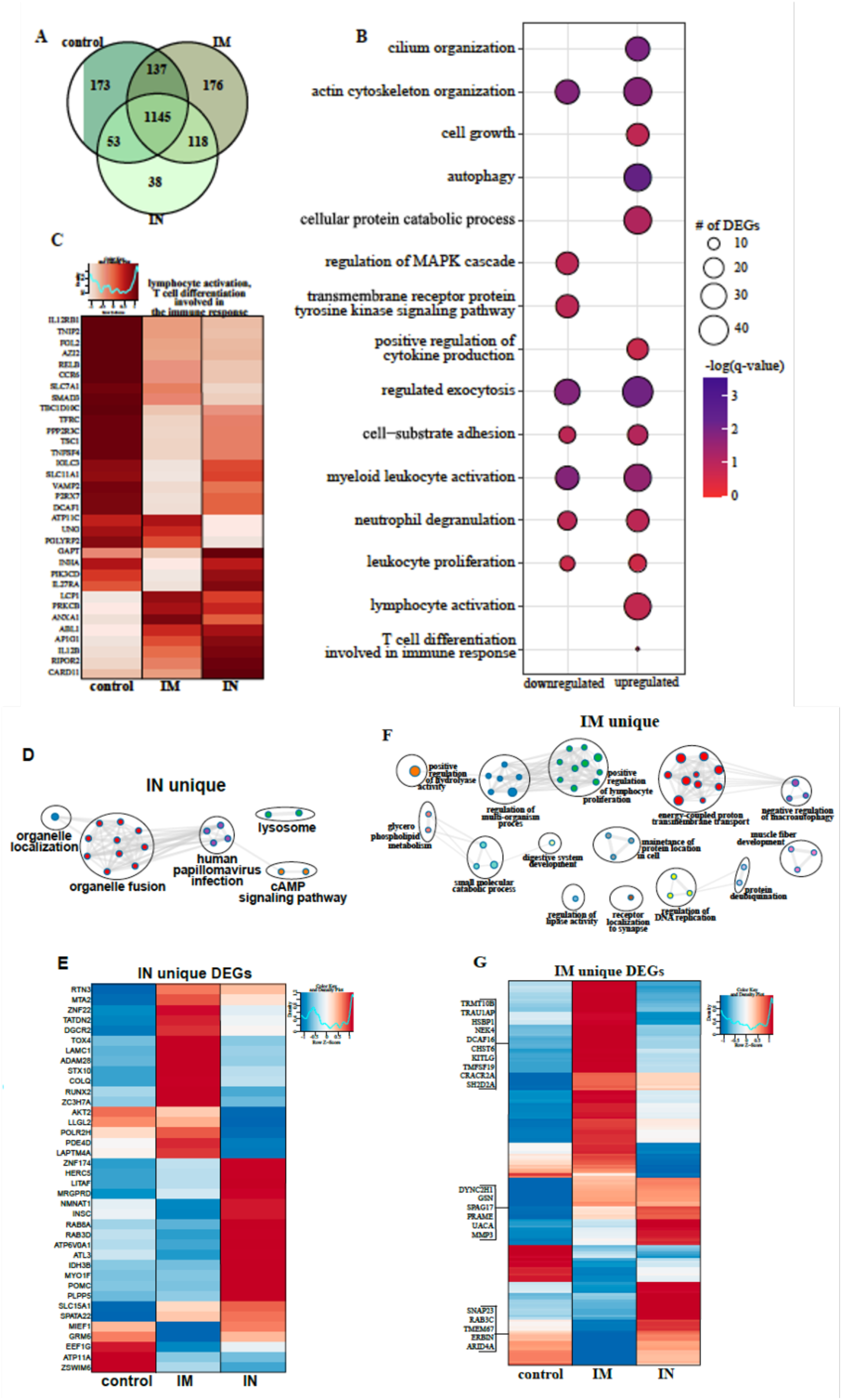
BAL RNA-sequencing. **(A)** Venn diagram of differentially expressed genes (DEGs) expressed 3 days post challenge with SARS-CoV-2. Animals either received a control, intramuscular (IM) or intranasal (IN) vaccination. **(B)** Bubbleplot representing functional enrichment of DEGs shared by all infected groups at 3 days post challenge. Color intensity of each bubble represents the negative log of p-value and the relative size of each bubble represents the number of DEGs belonging to the specified Gene Ontology (GO) term. **(C)** Heatmap representing shared upregulated DEGs enriching to GO terms “lymphocyte activation” and “T cell differentiation in volved in the immune response.” Expression is represented as the normalized rpkm, where each column represents the median rpkm of the given group. Range of colors is based on scale and centered rpkm values of the represented DEGs. GO term network depicting functional enrichment of DEGs unique to **(D)** IN and **(F)** IM using Mediascape. Color-coded clustered nodes correspond to one GO term or KEGG pathway. Node size represents the number of DEGs associated with the indicated term or pathway. Gray lines represent shared interactions between terms/pathways, with density and number indicating the strength of connections between closely related terms/pathways. Heatmaps representing DEGs unique to **(E)** IN and **(G)** IM. Exemplar DEGs are annotated. Red represents upregulation, blue presents downregulation. Each column represents the median rpkm of the given group. For all heatmaps, range of colors is based on scale and centered rpkm values of the represented DEGs.

Further analysis of shared DEGs showed that genes involved in pro-inflammatory pathways (e.g., *RELB, MFHAS1, IL12RB1, TNFSF4, TRAF2, C5AR1*, and *TOLLIP*) were more highly expressed in the control group compared to IM and IN groups (Fig. S6C, D). A second cluster of inflammation-related genes were induced to a greater extent in the IN group (*IFT88, IL12B, CLEC9*, and *IL27RA*), which is in line with the greater inflammatory response observed in these animals. On the other hand, DEGs implicated in T and B cell-mediated immunity (e.g., *LCP1, PRKCB*) were induced to a greater extent in IM-vaccinated NHPs (Fig. 5C). Genes enriching to GO term “myeloid cell activation/neutrophil downregulation” were suppressed to a greater extent in the IM cohort, consistent with the lower inflammation profile observed in this group (Fig. S6E).

We next analyzed vaccine-specific transcriptional responses in the BAL samples of the IN and IM groups to elucidate unique molecular responses (Fig. 5D-G). DEGs unique to the IN group enriched to GO terms related to organelle dynamics, such as “organelle localization” as well as antiviral immunity (“human papillomavirus infection”) (Fig. 5D). Genes important for vesicular mobilization (*RAB8A* and *RABD3D*) as well as antiviral HERC5 were more highly expressed in IN-vaccinated NHPs whereas those associated with signaling (e.g., *AKT2, PDE4D*) were more down-regulated (Fig. 5E) compared to IM-vaccinated NHPs. DEGs uniquely upregulated in the IM group have roles in protein synthesis and folding (e.g., *TRMT10B, TRNAU1AP, HSPB1*), cell proliferation (e.g., *KITLF, TM4SF19*) and T cell activation (e.g., *CRACR2A, SH2D2A*) (Fig. 5G).

Due to limited sample availability, we were unable to perform phenotyping of immune cells in the BAL. Therefore, we performed *in silico* flow cytometry to infer changes in cell frequencies based on the transcriptional landscape (Fig. S6F). This analysis predicted significant increases in the levels of monocytes, NK cells and stimulated CD4 Th2 cells while frequencies of naЇve & plasma B cell, CD4^+^ & CD8^+^ T cells, stimulated dendritic cells and neutrophils were predicted to decline for all three challenged groups. On the other hand, plasma cells and monocytes were predicted to be induced to a lower magnitude in controls compared to IM and IN groups (Fig. S6F).

### Transcriptional analysis of lung samples

As described for BAL samples, we observed a clear distinction between lung samples of uninfected/naЇve compared to vaccinated and challenged NHPs with no clear separation between the three challenged groups, therefore, we employed the same strategy as described above for BAL (Fig. S7A). A robust transcriptional response to SARS-CoV-2 infection was evident in all three vaccinated groups compared to naЇve animals (Fig. S7B) with the majority of DEGs shared among the three groups (Fig. 6A). Downregulated DEGs shared by all three vaccinated groups enriched to GO terms primarily involved in innate immunity (e.g., “regulation of innate immune response”, “antigen processing and presentation”), cellular stress (e.g., “coagulation”, “response to decreased oxygen levels”, “wound healing”) and cell cycle (e.g., “regulation of cell cycle process”) (Fig. 6B). Genes playing roles in protein folding and turnover (*CALR* and *CTSF*), immune activation (e.g., *LYN* and *ADA*) coagulation (e.g., *PLAT, SIRT2, FERMT3*), fluid homeostasis (e.g., *ADM, SERPINA5*) and cell morphogenesis (e.g., *NOTCH4, TEK*) were more suppressed in the IN group (Fig. 6C and S7C). Shared upregulated DEGs enriched to GO terms reflecting innate immune processes (e.g., “regulated exocytosis”, “myeloid leukocyte activation”, “neutrophil degranulation”), as well as cell migration (e.g., “chemotaxis”) and extracellular structural dynamics (e.g., “extracellular structural organization”, “cell projection morphogenesis”) (Fig. 6B). Overall, a large portion of these genes were induced to a greater extent in the IN group, notably those that play a role in neutrophil activation (e.g., *GRN, AZU1, CR1*), cell metabolism (e.g., *ALDOC, PGM1*, IMPDH1), chemotaxis (e.g., *CCL20, CCL8, CCL13, ICAM3*), and extracellular matrix remodeling (e.g., *MMP25, MMP16*) (Fig. 6D, S7D). Additionally, genes important for angiogenesis and apoptosis were more upregulated in the IN group (e.g., *VEGFD, PRKD1, SHC1, DAB2IP*) (Fig. S7E).

**Figure 6.**
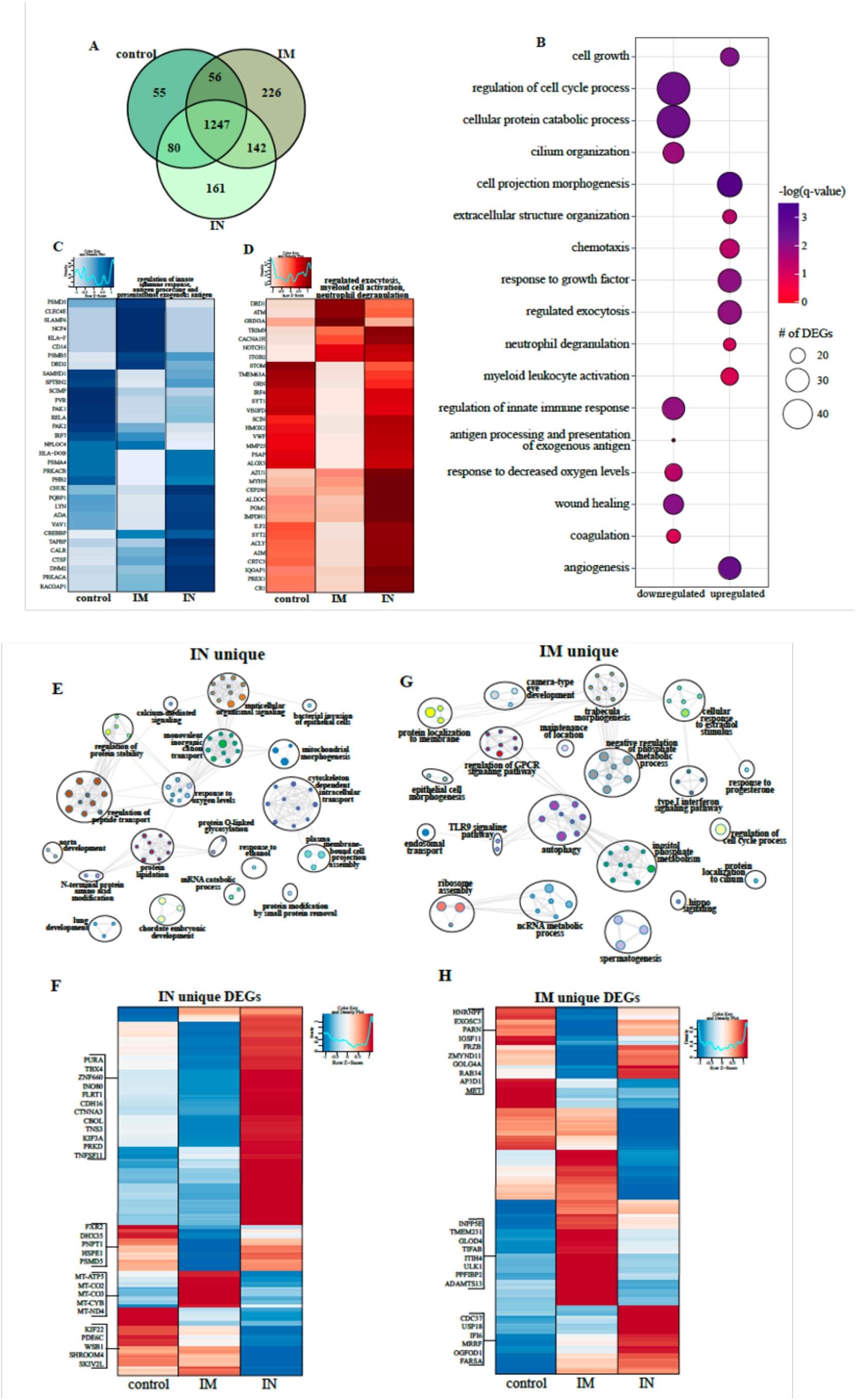
Lung RNA-Sequencing. **(A)** Venn diagram of differentially expressed genes (DEGs) expressed 3 days post challenge with SARS-CoV-2. Animals either received a control, intramuscular (IM) or intranasal (IN) vaccination. **(B)** Bubbleplot representing functional enrichment of DEGs shared by all infected groups at 7 days post challenge. Color intensity of each bubble represents the negative log of p-value and the relative size of each bubble represents the number of DEGs belonging to the specified Gene Ontology (GO) term. Heatmaps representing shared GO terms **(C)** “regulation of innate immune response”, “antigen processing and presentation of exogenous antigen” for downregulated DEGs; and **(D)** “regulated exocytosis”, “myeloid leukocyte activation” and “neutrophil degranulation” for upregulated DEGs. Expression is represented as the normalized rpkm, where each column represents the median rpkm of the given group. Range of colors is based on scale and centered rpkm values of the represented DEGs. GO term network depicting functional enrichment of DEGs unique to **(E)** IN and **(G)** IM using Mediascape. Color-coded clustered nodes correspond to one GO term or KEGG pathway. Node size represents the number of DEGs associated with the indicated term or pathway. Gray lines represent shared interactions between terms/pathways, with density and number indicating the strength of connections between closely related terms/pathways. Heatmaps representing DEGs unique to **(F)** IN and **(H)** IM. Exemplar DEGs are annotated. Red represents upregulation, blue presents downregulation. Each column represents the median rpkm of the given group. For all heatmaps, range of colors is based on scale and centered rpkm values of the represented DEGs.

We next analyzed vaccine route-specific DEGs to identify differences in the molecular responses to vaccination and challenge (Fig. 6E-H). Enrichment of the 161 DEGs unique to the IN group revealed enrichment to GO terms suggesting tissue injury (e.g., “lung development”), metabolism (e.g., “protein modification by small protein removal”), and signaling (e.g., “calcium-mediated signaling”) (Fig. 6F). Notably, downregulated DEGs in this group include components of the mitochondrial cellular respiration complex (e.g., *MT-CYB, MT-CO2*) and cellular homeostasis (e.g., *SKIV2L, PDE6C, SKIV2L, SHROOM4*) (Fig. 6F).

In contrast, the 226 DEGs unique to the IM group enriched to GO terms related to cellular defense (e.g., type I interferon signaling pathway”, “TLR9 signaling”, “autophagy”), morphogenesis (e.g., “epithelial cell morphogenesis”) and membrane dynamics (e.g., “protein localization to membrane”,) (Fig. 6G). The higher expression of DEGs related to the type I interferon response (e.g., *CDC37, USP18, IFI6*) and ribosome assembly (e.g., *MRRF, OGFOD1, FARSA*) in the IM and control groups suggests greater mobilization of host defense processes (Fig. 6H). Additionally, genes associated with cilia formation (e.g., *INPP5E, TMEM231, PPFIBP2*) and regulation of inflammation (e.g., *GLOD5, ITIH5, TIFAB*) were highly expressed in the IM group in line with reduced damage (Fig. 6H). *In silico* flow cytometry analysis indicated that these transcriptional changes were consistent with increases in the levels of neutrophils, monocytes, NK cells, dendritic cells and naЇve lymphocytes in all the infected animals while frequencies of activated monocytes were predicted to decrease relative to tissue from naЇve animals (Fig. S7F).

## Discussion

Many vaccine platforms have been utilized to develop a COVID-19 vaccine quickly (*13-16*), with several already approved for human use within a year of SARS-CoV-2 emergence. However, many of these vaccines require 2 doses to elicit protection and are delivered IM rather than the site of infection. Therefore, we developed a single dose, fast-acting VSV-based vaccine against COVID-19, which is based on the rVSV-ZEBOV vaccine approved by the US Food and Drug administration (FDA) and the European Medicines Agency (EMA) for human use. Additionally, we compared the protective efficacy of IM and IN delivery in the rhesus macaque model (*27*).

A single dose IM-, but not IN-delivered vaccine protected NHPs from COVID-19 pneumonia within 10 days post vaccination. This short time to immunity is a tremendous advantage and highlights its potential to be used rapidly during a public health crisis, particularly in emergency situations when many people were exposed at once. Interestingly, IM vaccination resulted in superior immune response compared to IN as evidenced by the significantly higher SARS-CoV-2-S-specific antibody titers and lower viral loads in this group. None of the animals in this study showed overt clinical signs of disease regardless of their vaccination status. However, histological examination of the lung tissue revealed immunopathology that was most significant in IN-vaccinated animals. The observed immunopathology was not consistent with a classic hypersensitivity response or immune-enhanced disease as the lesions were limited to the periphery of lung lobes and almost exclusively observed in lower lung lobes as previously reported for SARS-CoV-2 infection (*27*). Importantly, IM-vaccinated animals did not develop signs of interstitial pneumonia, nor could we detect SARS-CoV-2 antigen in the lungs. Indeed, at the time of euthanasia, lung lesions were apparent in animals from the IN and controls groups, but not in the IM group. This is surprising given mucosal vaccination for other respiratory pathogens has been demonstrated to be superior or comparable to IM (*30, 31*). It possible that the immunopathology observed in the vaccinated animals is due to the short duration between vaccination and challenge (10 days) and that it might also occur with other SARS-CoV-2 vaccine platforms, as this change was clinically silent in our model.

In line with the observations above, the transcriptional analysis of lung samples showed a divergence of antiviral states between the IN, IM and control groups. IN vaccination induced transcriptional changes enriched to cell metabolism, apoptosis, angiogenesis, and neutrophil activation processes. Some notable DEGs include genes associated with neutrophil activation and the formation of azurophil granules (e.g. *GRN, CR1*, and *AZU1*). While neutrophils play a role in protecting the host, sustained neutrophil activation has been shown to directly correlate to more severe COVID-19 cases (*32*). In contrast, IM vaccination induced transcriptional changes playing a role in cilia formation, inflammation regulation, and type I IFN. A majority of these genes are responsible for the regulation and control of early innate inflammation such as *USP18* which disrupts the JAK-STAT pathway downstream of the IFN receptor (*33*), and *TIFAB* which inhibits the activation of the NFkB pathway (*34*). Collectively these findings support the significant decrease in virus replication between the IN and IM vaccination groups. Transcriptional analysis of acute BAL samples also demonstrated a divergence of antiviral states evidenced by T cell differentiation genes upregulated after IM vaccination compared to innate antiviral posttranslational modifications after IN vaccination. While variable immunopathology was observed in all VSV-SARS2-EBOV-vaccinated NHPs regardless of the route, transcriptional analysis of the BAL demonstrated an upregulation of antiviral genes such as *HERC5* in the IN group only, in line with enhanced viral loads in this group. *HERC5* is responsible for the production of *HECT*-type E3 protein ligase, a facilitator of the ISG conjugation system of interferon stimulated gene 15 (*35*) possibly contributing to the enhanced immunopathology observed in the IN group.

Since the importance of a cellular immune response has been recently highlighted in COVID-19 patients (*36*), we assessed development of both innate as well as the adaptive cellular responses following each vaccination strategy. Our analysis showed a higher frequency of granzyme B^+^ NK cells after IM vaccination. While there was minimal antigen-specific cytokine production from the CD4^+^ T cells, a significant increase in the early activation marker, CD69, was observed on 0 and 7 dpc. In addition, an increase in the IL-2 production was only observed 7 dpc, indicative of the priming of cytotoxic CD8^+^ T cells (*37*). Transcriptional analysis revealed that IM vaccination induced upregulation of *CRACR2A* and *SH2D2a. SH2D2a* encodes a T cell-specific adaptor protein, which facilitates the formation and maintenance of the immunological synapse between the antigen presenting cell and the T cell receptor allowing for a more robust antigen-specific stimulation (*38*). *CRACR2A* also has a role in the maintenance of the immunological synapse and promotes downstream signaling, which results in an increased Th1 response and Th17 effector functions which is supported by the significant decrease of IL-2 on 0 dpc (*39*).

Serum cytokine analyses demonstrated a significant increase in IL-18 and innate chemokines, MCP-1 and MIP-1β, circulating in animals that presented with severe pathology. The increased expression of IL-18 could indicate a priming of the infiltrating immune cells in the lungs to a more proinflammatory state that would result in the observed tissue destruction. However, we did not observe an increase of IL-18 in BAL samples from 3 dpc. The most striking observation was the significant downregulation of MIP-1β on all dpc measured in IM-vaccinated animals. MIP-1β and MCP-1 have previously been demonstrated to be indicators of severe COVID-19 pathogenesis by transcriptomic profiling of human patients (*40*). A decrease in MIP-1β and MCP-1 could contribute to the lack of immune cell infiltration in the lungs of the IM group. Predicted *in silico* flow cytometry data from BAL on 3 dpc showed a decrease of naЇve and plasma B cells, CD4^+^ and CD8^+^ T cells, stimulated dendritic cells and neutrophils supporting this hypothesis. Furthermore, *in silico* flow cytometry from the lungs on 7 dpc were indicative of a decrease in activated monocytes for all SARS-CoV-2-infected NHPs. Further immune cell characterization from BAL and within the lungs is needed to expand upon our results and confirm this hypothesis.

In summary, in this study we generated a potent single-dose, fast-acting vaccine for COVID-19. This vaccine grows to high titers like the parent rVSV-ZEBOV vector and to higher titers compared to a VSV vaccine expressing the SARS-CoV-2 S alone (*41*)(unpublished data). Several important questions remain to be addressed in future studies. An extension of the time between vaccination and challenge might overcome the observed difference in protection between the vaccination routes and might eliminate the signs of immunopathology. This aspect will be investigated in future studies in conjunction with assessing the durability of SARS-CoV-2-specific immunity and a possible dose reduction of the vaccine as has been described for the rVSV-ZEBOV, the parental vaccine (*42*). The rVSV-ZEBOV has been shown to elicit a durable humoral response, which lasts for at least 2 years in humans (*43*). We will also investigate the addition of another SARS-CoV-2 antigen into the vaccine to promote a stronger T cell response as these responses are typically longer lasting. Furthermore, we will analyze if pre-existing immunity to EBOV could impact the immunogenicity of this bivalent vaccine. For now, the VSV-SARS2-EBOV vaccine presents a vaccine with a high potential as a boosting option after the already approved mRNA-based vaccine because the VSV-SARS2-EBOV elicits primarily a humoral response in contrast to the predominantly T cell-driven immune response after mRNA vaccination (*16*).

## Materials and Methods

### Ethics statement

All infectious work with SARS-CoV-2 was performed in the high containment laboratories at the Rocky Mountain Laboratories (RML), Division of Intramural Research, National Institute of Allergy and Infectious Diseases, National Institutes of Health. RML is an institution accredited by the Association for Assessment and Accreditation of Laboratory Animal Care International (AAALAC). All procedures followed standard operating procedures (SOPs) approved by the RML Institutional Biosafety Committee (IBC). Animal work was performed in strict accordance with the recommendations described in the Guide for the Care and Use of Laboratory Animals of the National Institute of Health, the Office of Animal Welfare and the Animal Welfare Act, United States Department of Agriculture. The studies were approved by the RML Animal Care and Use Committee (ACUC). Procedures were conducted in animals anesthetized by trained personnel under the supervision of veterinary staff. All efforts were made to ameliorate animal welfare and minimize animal suffering in accordance with the Weatherall report on the use of nonhuman primates in research (https://royalsociety.org/policy/publications/2006/weatherall-report/). Animals were housed in adjoining individual primate cages that enabled social interactions, under controlled conditions of humidity, temperature, and light (12 hours light - dark cycles). Food and water were available *ad libitum*. Animals were monitored and fed commercial monkey chow, treats, and fruit at least twice a day by trained personnel. Environmental enrichment consisted of commercial toys, music, video and social interaction.

### Animal study

Sixteen female rhesus macaques (3.5-10 years of age; 4.5-10kg, Indian-origin) were used in this study. The NHPs were randomly selected for two vaccine groups (n=6) and one control group (n=4). On day-10 NHPs received a single vaccine dose of 1×10^7^ PFU of VSV-SARS2-EBOV by the IM (injection caudal thigh) or IN route (dropping vaccine into each nostril). Control animals received the same dose of a control vaccine (VSV-EBOV) by the IM (n=2) or IN (n=2) route (Fig. S1D). On day 0, animals were challenged with SARS-CoV-2 as previously described (*27*). On day 0, 1, 3, 5 and 7 after challenge a clinical exam was performed including thoracic radiograph and nasal swab collection. The day 3 exam included bronchoalveolar lavage (BAL) using 10 ml sterile saline. On day 7, all animals were euthanized for sample collection.

### Cells and Viruses

Huh7 and VeroE6 cells were grown at 37°C and 5% CO_2_ in Dulbecco’s modified Eagle’s medium (DMEM) (Sigma-Aldrich, St. Louis, MO) containing 10% fetal bovine serum (FBS) (Wisent Inc., St. Bruno, Canada), 2 mM L-glutamine (Thermo Fisher Scientific, Waltham, MA), 50 U/mL penicillin (Thermo Fisher Scientific), and 50 μg/mL streptomycin (Thermo Fisher Scientific). BHK-T7 (baby hamster kidney) cells expressing T7 polymerase were grown at 37°C and 5% CO_2_ in minimum essential medium (MEM) (Thermo Fisher Scientific) containing 10% tryptose phosphate broth (Thermo Fisher Scientific), 5% FBS, 2 mM L-glutamine, 50 U/mL penicillin, and 50 μg/mL streptomycin. SARS-CoV-2 isolate nCoV-WA1-2020 (MN985325.1) (*44*) was used for the animal challenge studies and neutralization test.

### Generation of VSV-based vaccine candidates

The SARS-CoV-2 S ORF was PCR-amplified from an expression plasmid encoding the codon-optimized (human) gene based on GenBank accession number MN908947 which was kindly provided by Vincent Munster (NIAID). Full-length SARS-CoV-2 S was cloned into the pATX-VSV-EBOV plasmid upstream of the EBOV-Kikwit GP resulting in VSV-SARS2-EBOV (Fig. S1A) following a previously successful strategy (*45*). The replication-competent recombinant VSV was recovered in BHK-T7 cells as described previously (*26*). VSV-SARS2-EBOV was propagated on Huh7 cells. The complete sequence of the virus was confirmed by Sanger sequencing. The titer of the virus stock was quantified using standard plaque assay on VeroE6 cells.

### Growth kinetics

VeroE6 cells were grown to confluency in a 12-well plate and infected in triplicate with VSVwt, VSV-EBOV, or VSV-SARS2-EBOV at a multiplicity of infection of 0.01. After 1 h incubation at 37°C, cells were washed three times with plain DMEM, and covered with DMEM containing 2% FBS. Supernatant samples were collected at 0, 6, 12, 24, 48, 72, and 96 hours post infection and stored at −80°C. The titer of the supernatant samples was determined performing TCID_50_ assay on VeroE6 cells as previously described (*26*).

### Western blot analysis

Supernatant samples containing VSV were mixed 1:1 with sodium dodecyl sulfate-polyacrylamide (SDS) gel electrophoresis sample buffer containing 20% β-mercaptoethanol and heated to 99°C for 10min. SDS-PAGE and transfer to Trans-Blot polyvinylidene difluoride membranes (Bio-Rad Laboratories) of all samples was performed as described elsewhere (*22*). Protein detection was performed using anti-SARS-CoV-2 S RBD (1:1000; Sino Biological) or anti-EBOV GP (ZGP 12/1.1, 1μg/ml; kindly provided by Ayato Takada, Hokkaido University, Japan) or anti-VSV M (23H12, 1:1000; Kerafast Inc.). After horse-radish peroxidase (HRP)-labeled secondary antibody staining using either anti-mouse IgG (1:10,000) or anti-rabbit IgG (1:5000) (Jackson ImmunoResearch), the blots were imaged using the SuperSignal West Pico chemiluminescent substrate (Thermo Fisher Scientific) and an iBright™ CL1500 Imaging System (Thermo Fisher Scientific).

### RNA extraction and RT-qPCR

Blood, BAL fluid, and nasal swabs were extracted using the QIAamp Viral RNA Mini Kit (QIAGEN) according to manufacturer specifications. Tissues, a maximum of 30 mg each, were processed and extracted using the RNeasy Mini Kit (QIAGEN) according to manufacturer specifications. One step RT-qPCR for both genomic and subgenomic viral RNA was performed using specific primer-probe sets and the QuantiFast Probe RT-PCR +ROX Vial Kit (QIAGEN), in the Rotor-Gene Q (QIAGEN) as described previously (*13*). Five μL of each RNA extraction were run alongside dilutions of SARS-CoV-2 standards with a known concentration of RNA copies.

### Enzyme-linked immunosorbent assay

Serum and BAL samples from SARS-CoV-2-infected animals were inactivated by γ-irradiation and used in BSL2 according to IBC-approved SOPs. NUNC Maxisorp Immuno plates were coated with 50 μl of 1 μg/mL of recombinant SARS-CoV-2 spike (S1+S2), SARS-CoV-2 RBD (Sino Biological) or EBOV GP antigen at 4°C overnight and then washed three times with phosphate buffer saline containing 0.05% Tween 20 (PBST). The plates were blocked with 3% skim milk in PBS for 3 hours at room temperature, followed by three additional washes with PBST. The plates were incubated with 50 μl of serial dilutions of the samples in PBS containing 1% skim milk for 1hour at room temperature. After 3 washes with PBST, the bound antibodies were labeled using 50 μl of 1:2,500 horse-radish peroxidase (HRP)-labeled anti-monkey IgG (H+L) (SeraCare Life Sciences) diluted in 1% skim milk in PBST. For the IgG subclass ELISAs the plates were incubated with samples at 4°C overnight. After three washes with PBST, 50 μl of 1 μg/mL Anti-rhesus IgG1 [ena], IgG2 [dio], IgG3 [tria], or IgG4 [tessera] (NHPRR) diluted in 1% skim milk in PBST was added and incubated for 1 h at room temperature. After 3 washes with PBST, the bound antibodies were labeled using 50 μl of 1:10,000 HRP-labeled anti-mouse IgG (H+L) (SeraCare Life Sciences) diluted in 1% skim milk in PBST. For all ELISAs, after incubation for 1 h at room temperature and 3 washes with PBST, 50 μl of KPL ABTS peroxidase substrate solution mix (SeraCare Life Sciences) was added to each well, and the mixture was incubated for 30 min at room temperature. The optical density (OD) at 405 nm was measured using a GloMax® explorer (Promega). The OD values were normalized to the baseline samples obtained on day −10 and the cutoff value was set as the mean OD plus standard deviation of the blank.

### Virus neutralization assay

The day before this assay, VeroE6 cells were seeded in 96-well plates. Serum samples were heat-inactivated for 30 min at 56°C, and 2-fold serial dilutions were prepared in DMEM with 2% FBS. Next, 100 TCID_50_ of SARS-CoV-2 were added and the mixture was incubated for 1 hour at 37°C and 5% CO_2_. Finally, media was removed from cells and the mixture was added to VeroE6 cells and incubated at 37°C and 5% CO_2_ for 6 days. Then the CPE was documented, and the virus neutralization titer was expressed as the reciprocal value of the highest dilution of the serum which inhibited virus replication (no CPE).

### Flow cytometry

Rhesus macaque PBMCs were isolated from ethylene diamine tetraceticacid (EDTA) whole blood by overlay on a Histopaque®-1077 density cushion and separated according to manufacturers’ instructions. Isolated PBMCs were resuspended in FBS with 10% DMSO and frozen at −80°C until analysis. For analysis of T cell intracellular cytokine production, cells were stimulated for 6 hours with 1μg/ml SARS-CoV-S peptide pool, media, cell stimulation cocktail (containing PMA-Ionomycin, Biolegend), or Lassa virus (LASV) GP peptide pool together with 5μg/ml Brefeldin A (Biolegend). Following surface staining with Live/Dead-APCCy7, CD3-FITC, CD4-Alexa700, CD8-PeTexas Red, CD56-BV421 and CD69-PeCy7, cells were fixed with 4% paraformaldehyde (PFA) and stained intracellularly with IFNγ-BV605, IL-4-APC, IL-2-PerCPCy5.5 diluted in perm-wash buffer (Biolegend). For analysis of NK cell intracellular cytokine production, cells were stimulated as described above. Following surface staining with Live/Dead-APCCy7, CD3-FITC, CD4-PerCPCy5.5, CD8-PeTexas Red, CD16-Alexa700, and CD56-BV421, cells were fixed with 4% PFA and stained intracellularly with granzyme B-APC. Sample acquisition was performed on a Cytoflex-S (Beckman Coulter) and data analyzed in FlowJo V10 (TreeStar). Antigen specific T cells were identified by gating on Live/Dead negative, doublet negative (SSC-H vs SSC-A), CD3^+^, CD56^-^, CD4^+^ or CD8^+^ cells and cytokine positive. Three NK cell sub-populations were identified by gating on Live/Dead negative, doublet negative (SSC-H vs SSC-A), CD3^-^, CD56^-^, and CD8^+^ or CD16^+^ or CD8^+^CD16^+^ double positive. Cytokine responses for each sub-population were identified by gating on the population then granzyme B^+^ cells. Cytokine positive responses are presented after subtraction of the background responses detected in the LASV GP peptide stimulated samples.

### Cytokine analysis

Macaque serum and BAL samples were inactivated by γ-irradiation and removed from the high containment laboratory according to IBC-approved SOPs. Samples were then diluted 1:2 in serum matrix for analysis with Milliplex Non-Human Primate Magnetic Bead Panel as per manufacturer’s instructions (Millipore Corporation). Concentrations for each cytokine were determined for all samples using the Bio-Plex 200 system (BioRad Laboratories Inc.).

### Histology and immunohistochemistry

Necropsies and tissue sampling were performed according to IBC-approved SOPs. Lungs were perfused with 10% formalin and processed for histologic review. Harvested tissues were fixed for eight days in 10% neutral-buffered formalin, embedded in paraffin, processed using a VIP-6 Tissue Tek (Sakura Finetek, USA) tissue processor, and embedded in Ultraffin paraffin polymer (Cancer Diagnostics, Durham, NC). Samples were sectioned at 5 μm, dried overnight at 42 °C, and resulting slides were stained with hematoxylin and eosin. Specific anti-CoV immunoreactivity was detected using an in-house SARS-CoV-2 nucleocapsid protein (U864YFA140-4/CB2093) rabbit antibody (Genscript) at a 1:1000 dilution. The IHC assay was carried out on a Discovery ULTRA automated staining instrument (Roche Tissue Diagnostics) with a Discovery ChromoMap DAB (Ventana Medical Systems) kit. All tissue slides were evaluated by a board-certified veterinary pathologist. Sections taken at 3 levels from each lung lobe, totally 18 sections, were evaluated for each animal; a representative lesion from each group was selected for Fig. 3.

### Library construction and sequencing

Quality and quantity of RNA from BAL and lower left lung (LLL) were determined using an Agilent 2100 Bioanalyzer. cDNA libraries were constructed using the NEB Next Ultra II Direction RNA Library Prep Kit (Thermo Fischer). RNA was treated with RNase H and DNase I following depletion of ribosomal RNA (rRNA). Adapters were ligated to cDNA products and the subsequent ∼300 base pair (bp) amplicons were PCR-amplified and selected by size exclusion. cDNA libraries were assessed for quality and quantity prior to 150 bp single-end sequencing using the Illumina NovaSeq platform.

### Bioinformatic analysis

Preliminary data analysis was performed with RNA-Seq workflow module of systemPipeR, developed by Backman *et. al* (*46*). RNA-Seq reads were demultiplexed, quality-filtered and trimmed using Trim Galore (average Phred score cut-off of 30, minimum length of 50 bp). FastQC was used to generate quality reports. Hisat2 was used to align reads to the reference genome *Macaca mulatta* (Macaca_mulatta.Mmul_8.0.1.dna.toplevel.fa) and the Macaca_mulatta.Mmul_8.0.1.97.gtf was used for annotation. For viral read quantification, RNA-Seq reads were separately aligned to the severe acute respiratory syndrome coronavirus 2 Wuhan isolate genome (NC_045512.2) and the GCF_009858895.2_ASM985889v3_genomic.gff annotation file was used. Raw expression values (gene-level read counts) were generated using the summarizeOverlaps function and normalized (read per kilobase of transcript per million mapped reads, rpkm) using the edgeR package. Statistical analysis with edgeR was used to determine differentially expressed genes (DEGs) meeting the following criteria: genes with median rpkm of ≥5, a false discovery rate (FDR) corrected p-value ≤ 0.05 and a log_2_fold change ≥ 1 compared to uninfected tissues. The number of total viral reads was determined as the total number of normalized read counts mapping to all viral genes.

Functional enrichment of DEGs was performed using Metascape to identify relevant Gene Ontology (GO) biological process terms and KEGG pathways. *In silico* flow cytometry was performed using ImmQuant with the IRIS database. Heatmaps, bubbleplots, Venn diagrams and violin plots were generated using R packages ggplot2 and VennDiagrams. GO network plots were generated in Cytoscape (Version 3.5.1). Graphs were generated using GraphPad Prism software (version 8).

### Statistical analyses

All statistical analysis was performed in Prism 8 (GraphPad). The *in vitro* growth kinetics of recombinant VSVs (Fig. S1C) was examined using two-way ANOVA with Tukey’s multiple comparisons to evaluate statistical significance at all timepoints. Bioinformatics data were analyzed using one-way ANOVA with multiple comparisons, comparisons were made to either uninfected animals or control-vaccinated animals. Two-tailed Mann-Whitney’s rank or Wilcoxon tests were conducted to compare differences between groups for all other data. A Bonferroni correction was used to control for type I error rate where required. Statistically significant differences are indicated as follows: p<0.0001 (****), p<0.001 (***), p<0.01 (**) and p<0.05 (*).

## Acknowledgements

We thank the Rocky Mountain Veterinary Branch, NIAID for supporting the animal studies, and Anita Mora (NIAID) for assistance generating the pathology figures.

## Funding

The study was funded by the Intramural Research Program, NIAID, NIH. RNA sequencing was funded by the National Center for Research Resources and the National Center for Advancing Translational Sciences, NIH, through grant UL1 TR001414 awarded to I.M.

## Author contributions

A.M. conceived the idea and secured funding. W.F. and A.M. designed the studies. W.F., K.S., A.J.G., F.F., A.O., T.G., J.L., P.W.H., T.T., C.S.C., K.L.O., and A.M. conducted the studies, processed the samples and acquired the data. A.N.P. and A.J. performed the transcriptomics work. W.F., K.S., A.N.P., C.S.C., I.M., K.L.O., and A.M. analyzed and interpreted the data. W.F., I.M., K.L.O., and A.M. prepared the manuscript. All authors approved the manuscript.

## Competing interest

The authors declare no conflicts of interest.

## Data availability

All data is available in the manuscript or the supplementary materials.

---

**Figure S1.**
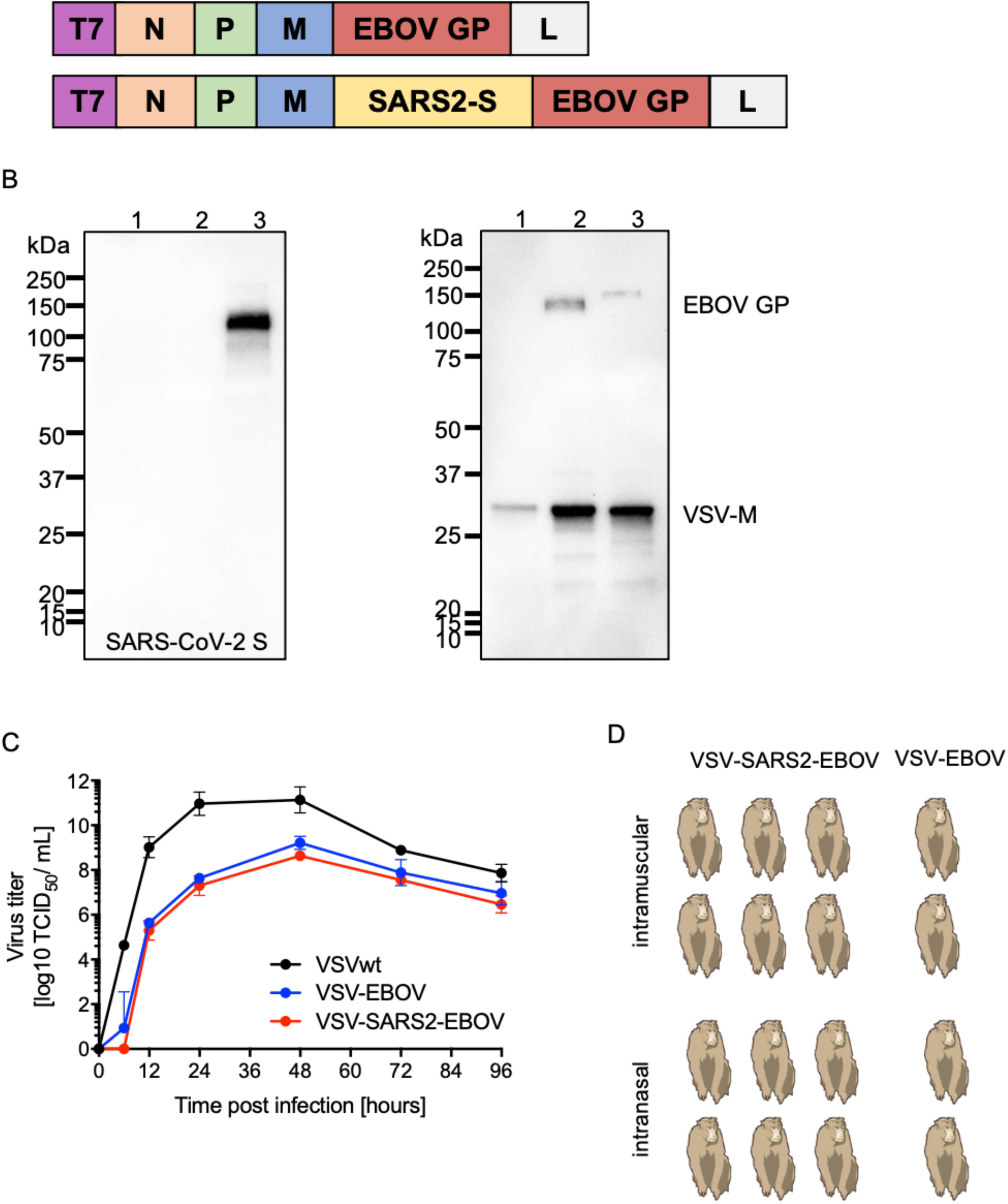
Schematic and characterization of VSV-based vaccines. **(A)** Schematic illustrating vaccine vector design. T7 promotor; N nucleoprotein; P phosphoprotein; M matrix protein; EBOV GP Ebola virus glycoprotein; L RNA-dependent RNA polymerase; SARS2-S SARS-CoV-2 S. **(B)** Western blot analysis of cell supernatant samples containing VSV vaccines probed for SARS-CoV-2 S (left), VSV M (middle) or EBOV GP (right). 1 VSV wildtype (VSVwt); 2 VSV-EBOV; 3 VSV-SARS2-EBOV. **(C)** Viral growth kinetics on VeroE6 cells. Geometric mean and SD are depicted. Results are not statistically significant. **(D)** Schematic outline of the rhesus macaque study.

**Figure S2.**
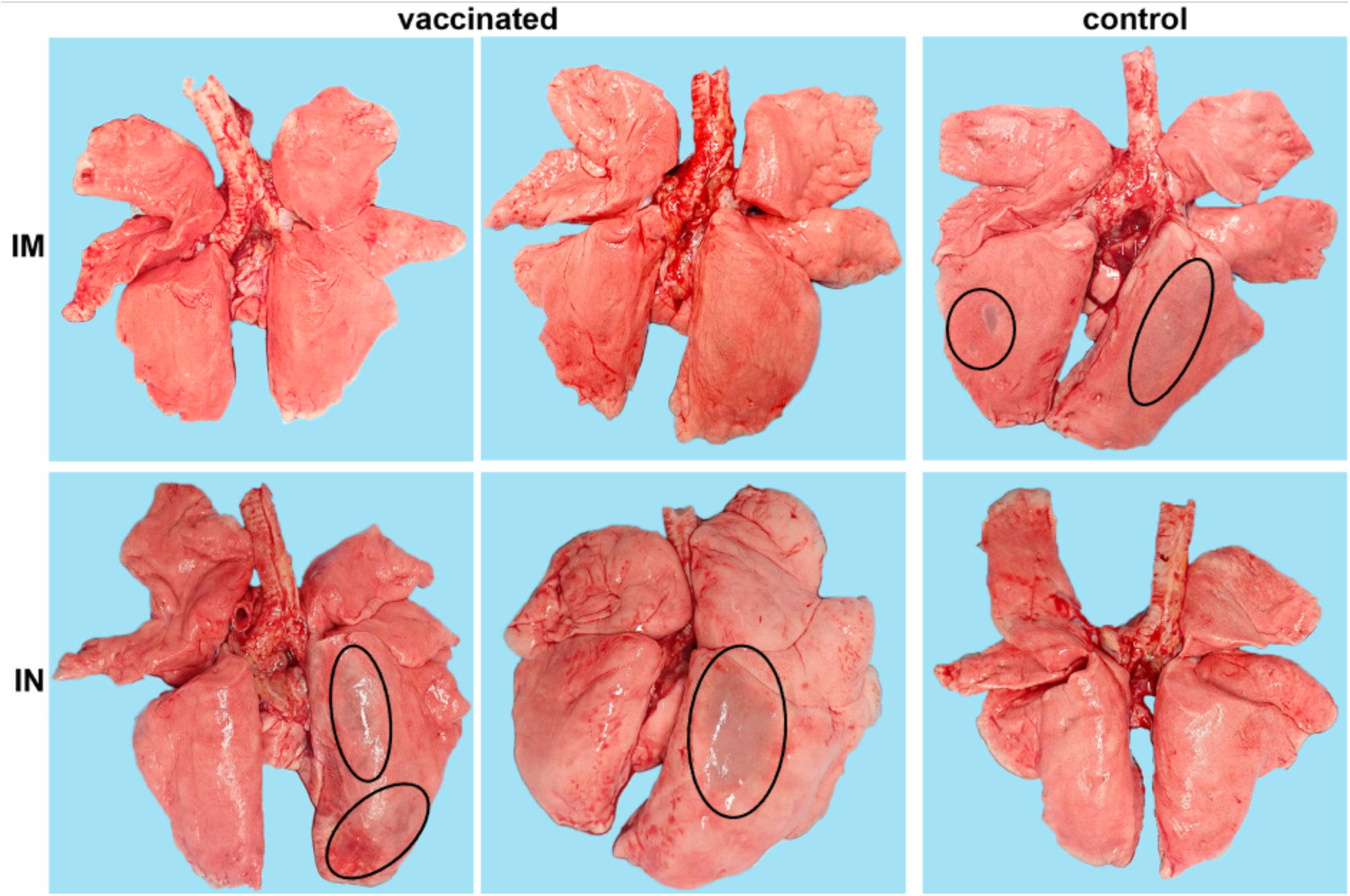
Lung gross pathology of NHPs. Representative pictures of NHP lungs with lesions on day 7 are shown. IM intramuscular vaccination; IN intranasal vaccination. Lesions are circled.

**Figure S3.**
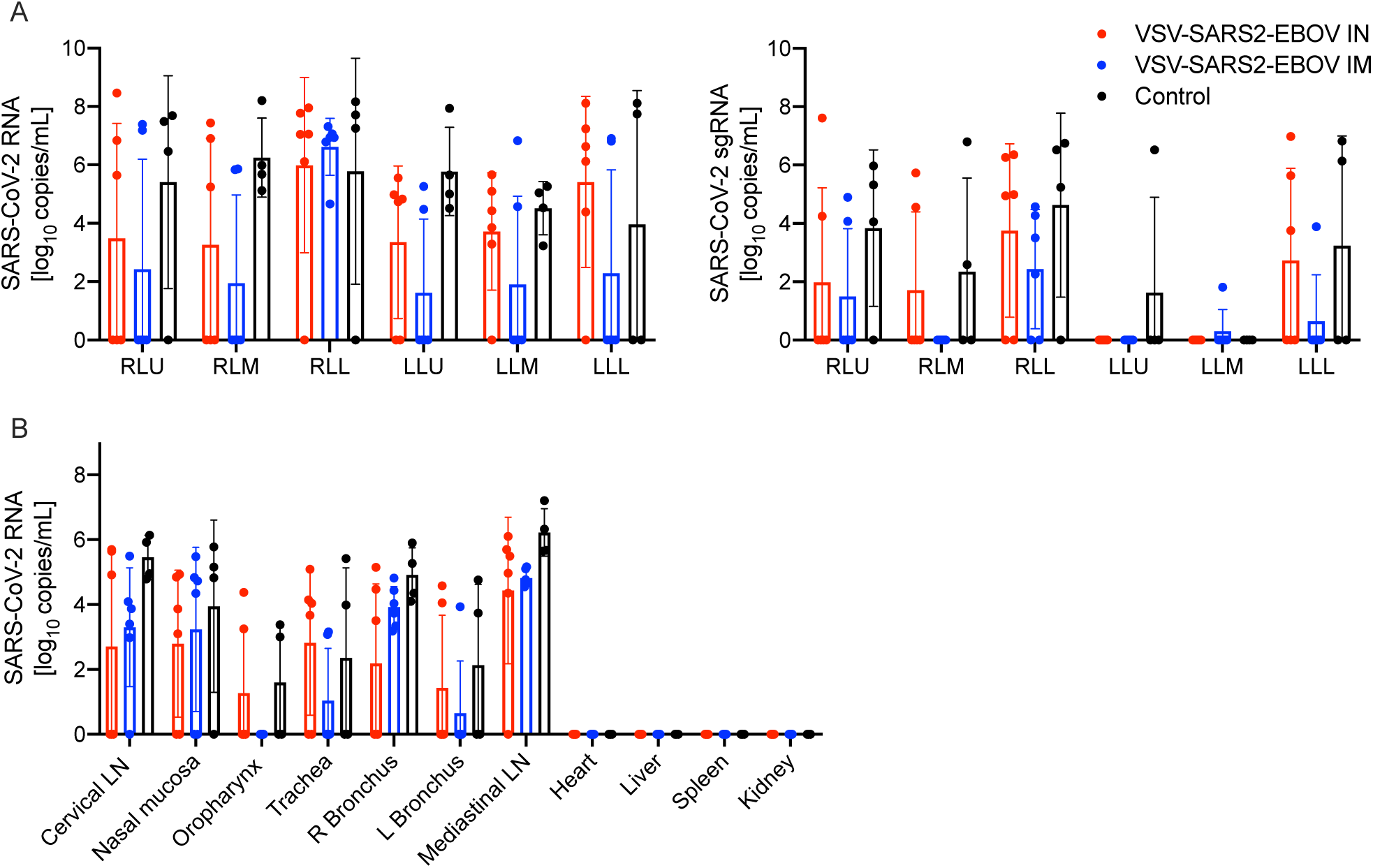
Virus load in NHP tissue samples on day 7. **(A)** Total SARS-CoV-2-specific RNA (left panel) and subgenomic (sg) RNA (right panel) in lung samples collected from NHPs. RLU right lobe upper; RLM right lobe middle; RLL right lobe lower; LLU left lobe upper; LLM left lobe middle; LLL left lobe lower. **(B)** Total SARS-CoV-2-specific RNA in tissue samples (right panel) from NHPs. LN lymph node; R right; L left.

**Figure S4.**
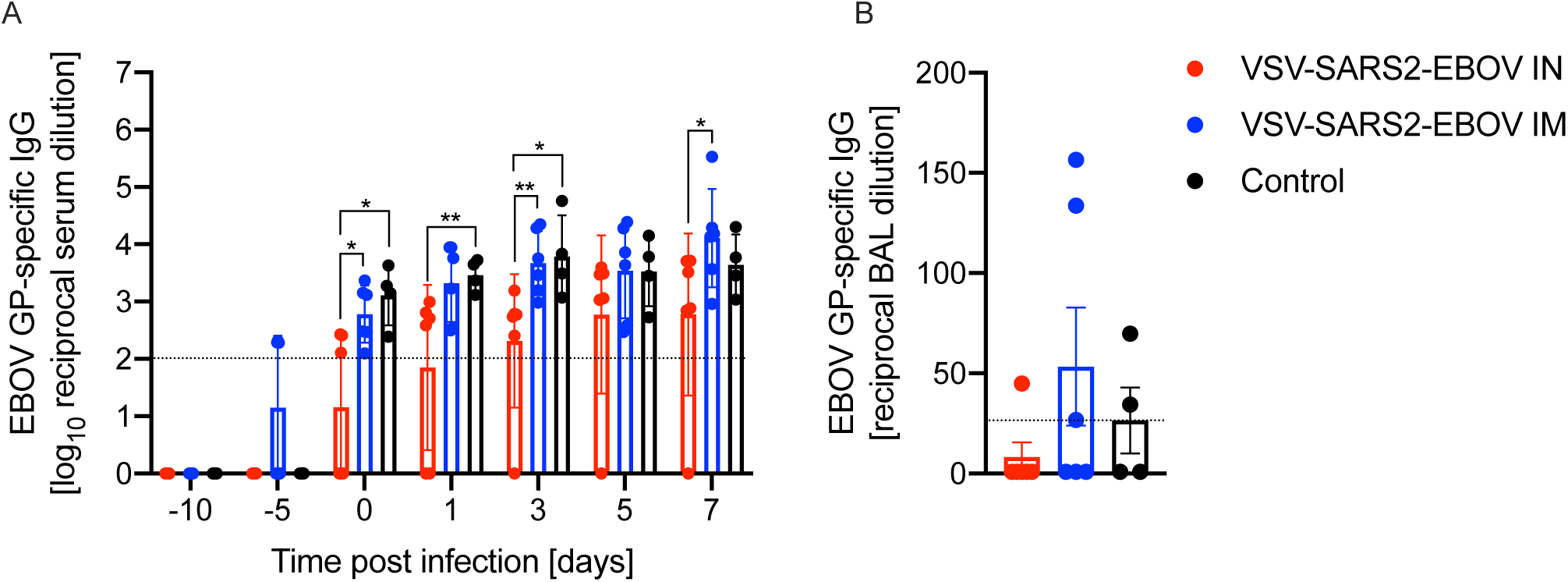
EBOV GP-specific antibodies in NHPs after vaccination and challenge. Bronchoaveolar lavage (BAL) samples collected on day 3 and serum samples collected throughout the study were analyzed by ELISA for EBOV GP-specific IgG. Statistical significance is indicated.

**Figure S5.**
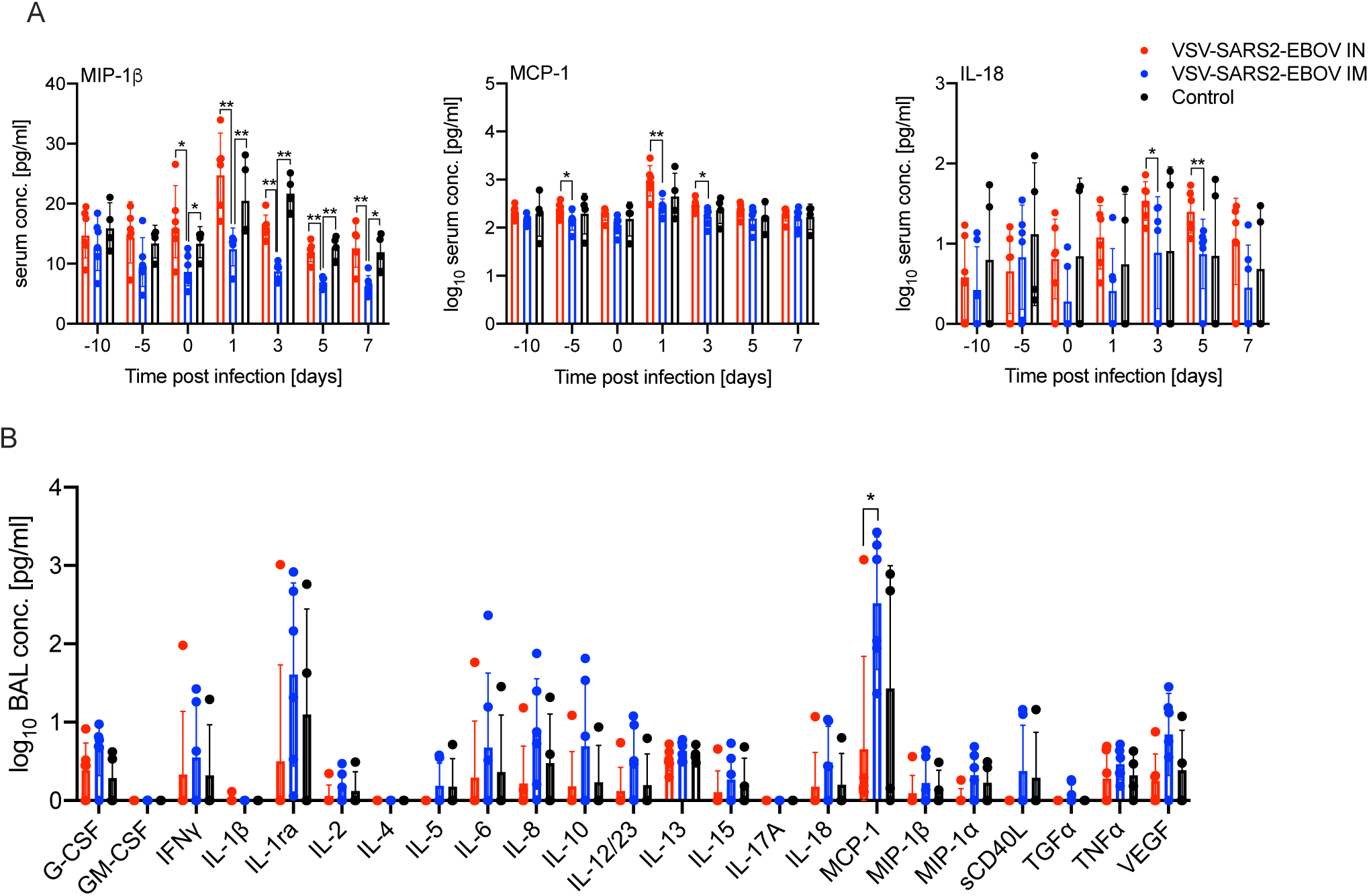
Serum cytokine levels in NHPs after vaccination and challenge. **(A)** Serum samples collected throughout the study and **(B)** bronchoaveolar lavage (BAL) samples collected on day 3 were analyzed. Statistical significance is indicated.

**Figure S6.**
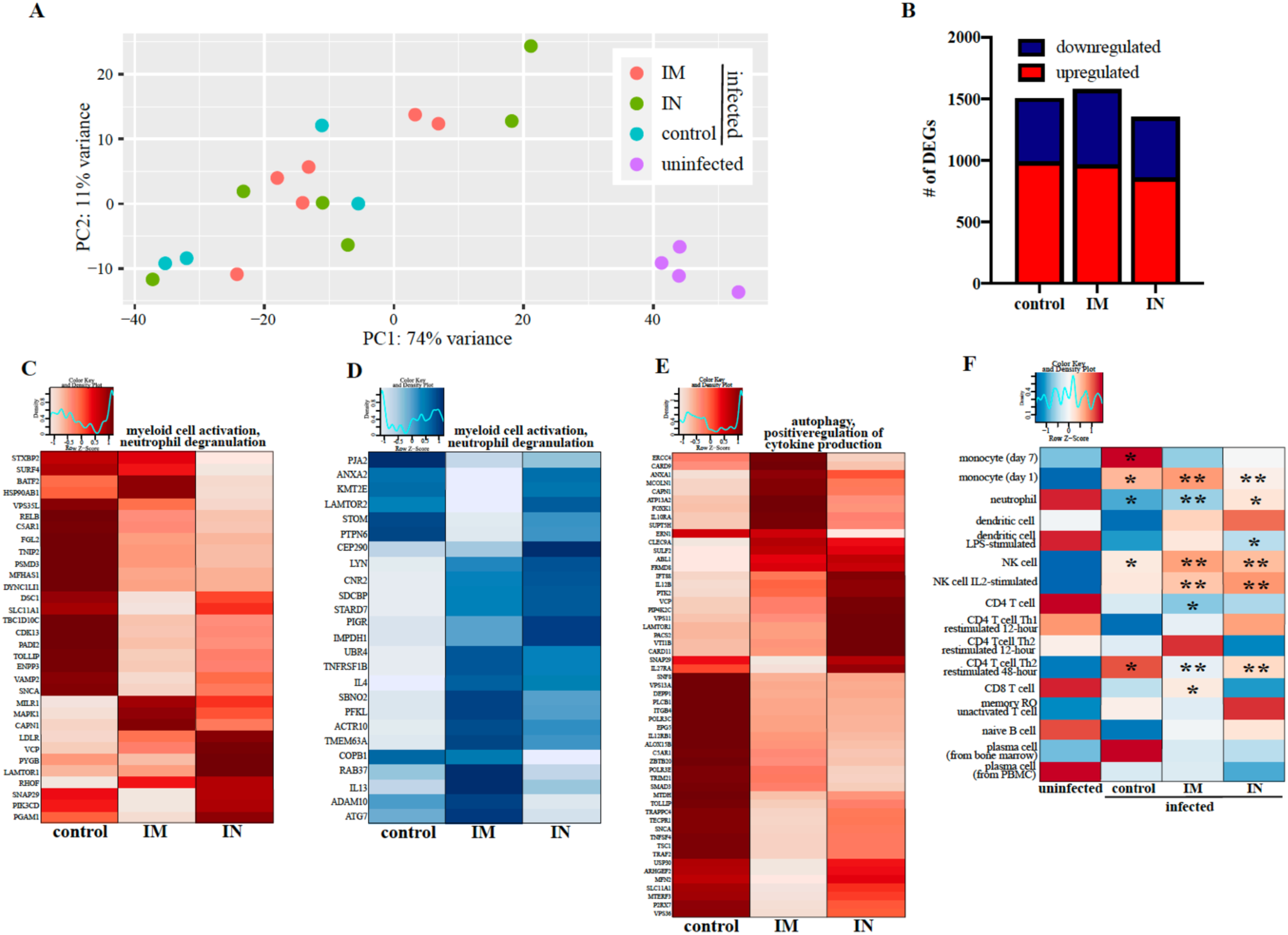
BAL RNA-sequencing. **(A)** Principal component analysis of bronchoalveolar lavage (BAL) samples from uninfected animals and vaccinated animals 3 days post challenge (control, intramuscular (IM) or intranasal (IN) vaccination). **(B)** Down- and up-regulated differentially expressed genes (DEGs). Heatmaps representing DEGs shared by all infected groups and enriching to Gene Ontology (GO) terms “myeloid cell activation” and “neutrophil degranulation” for **(C)** upregulated DEGs and **(D)** downregulated DEGs, and **(E)** “autophagy” and “positive regulation of cytokine production” for upregulated DEGs. Each column represents the median rpkm of the given group. Range of colors is based on scale and centered rpkm values of the represented DEGs. Red represents upregulated DEGs; blue represents downregulated DEGs. **(F)** *In silico* flow cytometry using ImmQuant IRIS database comparing challenged groups to uninfected controls. Red represents upregulation; blue represents downregulation. Each column represents the average relative predicted frequency of the given cell type. P-values are calculated relative to the uninfected animals. Statistical significance is indicated. For all heatmaps, range of colors is based on scale and centered rpkm values of the represented DEGs.

**Figure S7.**
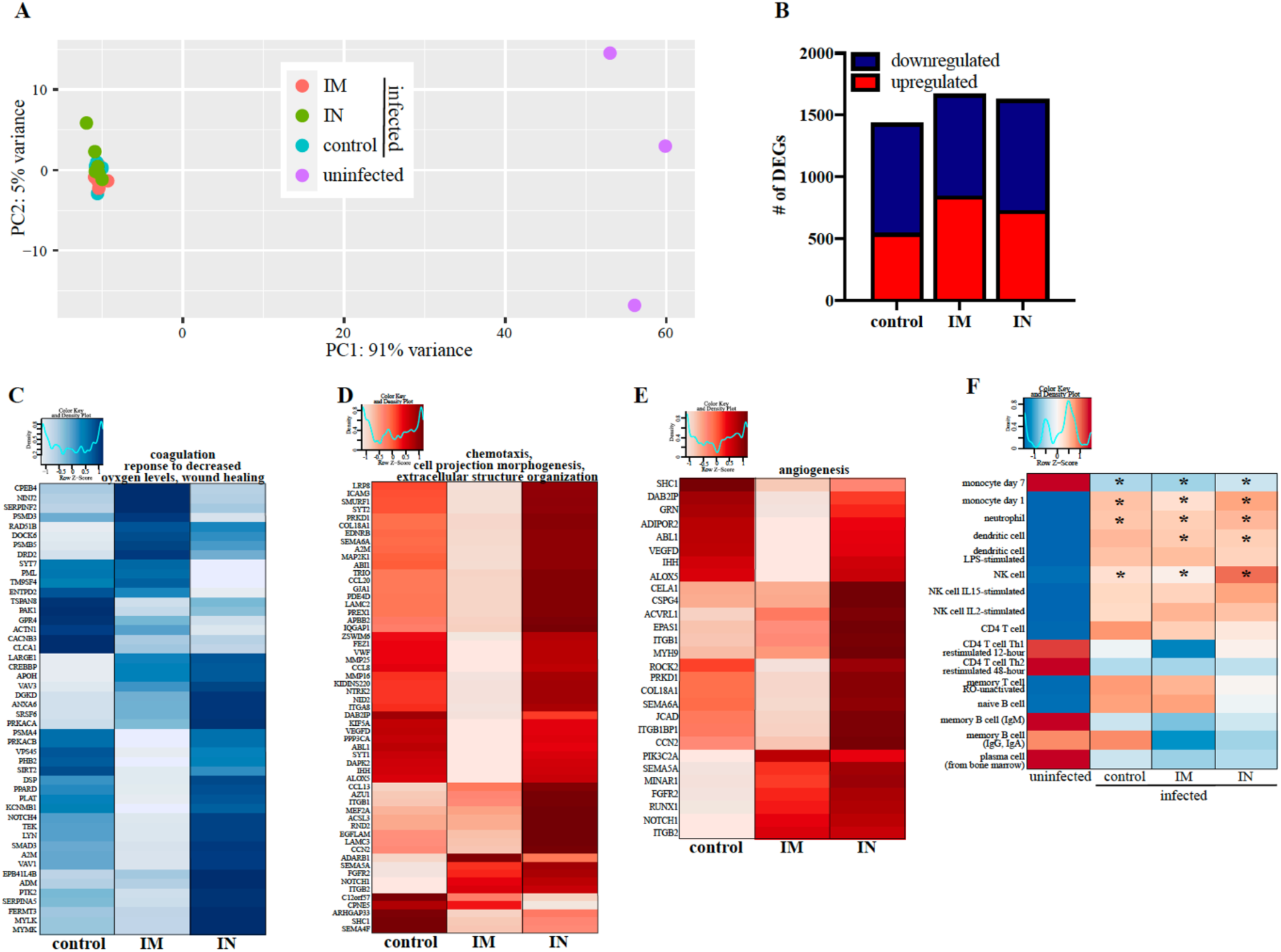
Lung RNA-sequencing. **(A)** Principal component analysis of lower left lung (LLL) samples from uninfected animals and vaccinated animals 7 days post challenge (control, intramuscular (IM) or intranasal (IN) vaccination). **(B)** Down- and up-regulated differentially expressed genes (DEGs). Heatmaps representing DEGs shared by all infected groups and enriching to Gene Ontology (GO) terms **(C)** “coagulation”, “response not decreased oxygen levels” and “wound healing” for downregulated DEG; and **(D)** “chemotaxis”, “cell projection morphogenesis” and “extracellular structure organization” and **(E)** “angiogenesis” for upregulated DEGs. Each column represents the median rpkm of the given group. Range of colors is based on scale and centered rpkm values of the represented DEGs. Red scale represents upregulated DEGs; blue scale represents downregulated DEGs. **(F)** *In silico* flow cytometry using ImmQuant IRIS database comparing challenged groups to uninfected controls. Red represents upregulation; blue represents downregulation. Each column represents the average relative predicted frequency of the given cell type. P-values are calculated relative to the uninfected animals. Statistical significance is indicated. For all heatmaps, range of colors is based on scale and centered rpkm values of the represented DEGs.

## Notes

### Competing Interest Statement

The authors have declared no competing interest.

## References

1. P. Zhou et al., A pneumonia outbreak associated with a new coronavirus of probable bat origin. Nature 579, 270–273 (2020).

2. W. J. Guan et al., Clinical Characteristics of Coronavirus Disease 2019 in China. N Engl J Med 382, 1708–1720 (2020).

3. E. W. Cheung et al., Multisystem Inflammatory Syndrome Related to COVID-19 in Previously Healthy Children and Adolescents in New York City. JAMA 324, 294–296 (2020).

4. R. Mao et al., Manifestations and prognosis of gastrointestinal and liver involvement in patients with COVID-19: a systematic review and meta-analysis. Lancet Gastroenterol Hepatol 5, 667–678 (2020).

5. D. Wichmann et al., Autopsy Findings and Venous Thromboembolism in Patients With COVID-19: A Prospective Cohort Study. Ann Intern Med 173, 268–277 (2020).

6. F. Zhou et al., Clinical course and risk factors for mortality of adult inpatients with COVID-19 in Wuhan, China: a retrospective cohort study. Lancet 395, 1054–1062 (2020).

7. F. Wu et al., A new coronavirus associated with human respiratory disease in China. Nature 579, 265–269 (2020).

8. M. L. Holshue et al., First Case of 2019 Novel Coronavirus in the United States. N Engl J Med 382, 929–936 (2020).

9. Q. Li et al., Early Transmission Dynamics in Wuhan, China, of Novel Coronavirus-Infected Pneumonia. N Engl J Med 382, 1199–1207 (2020).

10. M. Letko, A. Marzi, V. Munster, Functional assessment of cell entry and receptor usage for SARS-CoV-2 and other lineage B betacoronaviruses. Nat Microbiol 5, 562–569 (2020).

11. B. Ju et al., Human neutralizing antibodies elicited by SARS-CoV-2 infection. Nature 584, 115–119 (2020).

12. A. O. Hassan et al., A Single-Dose Intranasal ChAd Vaccine Protects Upper and Lower Respiratory Tracts against SARS-CoV-2. Cell 183, 169–184 e113 (2020).

13. N. van Doremalen et al., ChAdOx1 nCoV-19 vaccine prevents SARS-CoV-2 pneumonia in rhesus macaques. Nature 586, 578–582 (2020).

14. J. Yu et al., DNA vaccine protection against SARS-CoV-2 in rhesus macaques. Science 369, 806–811 (2020).

15. N. B. Mercado et al., Single-shot Ad26 vaccine protects against SARS-CoV-2 in rhesus macaques. Nature 586, 583–588 (2020).

16. K. S. Corbett et al., Evaluation of the mRNA-1273 Vaccine against SARS-CoV-2 in Nonhuman Primates. N Engl J Med 383, 1544–1555 (2020).

17. A. Marzi, H. Feldmann, T. W. Geisbert, D. Falzarano, Vesicular Stomatitis Virus-Based Vaccines for Prophylaxis and Treatment of Filovirus Infections. J Bioterror Biodef S1, (2011).

18. C. E. Mire et al., Use of Single-Injection Recombinant Vesicular Stomatitis Virus Vaccine to Protect Nonhuman Primates Against Lethal Nipah Virus Disease. Emerg Infect Dis 25, 1144–1152 (2019).

19. D. Safronetz et al., A recombinant vesicular stomatitis virus-based Lassa fever vaccine protects guinea pigs and macaques against challenge with geographically and genetically distinct Lassa viruses. PLoS Negl Trop Dis 9, e0003736 (2015).

20. A. Fathi, C. Dahlke, M. M. Addo, Recombinant vesicular stomatitis virus vector vaccines for WHO blueprint priority pathogens. Hum Vaccin Immunother 15, 2269–2285 (2019).

21. A. Marzi et al., EBOLA VACCINE. VSV-EBOV rapidly protects macaques against infection with the 2014/15 Ebola virus outbreak strain. Science 349, 739–742 (2015).

22. W. Furuyama et al., A single dose of a vesicular stomatitis virus-based influenza vaccine confers rapid protection against H5 viruses from different clades. NPJ Vaccines 5, 4 (2020).

23. K. S. Brown, D. Safronetz, A. Marzi, H. Ebihara, H. Feldmann, Vesicular stomatitis virus-based vaccine protects hamsters against lethal challenge with Andes virus. J Virol 85, 12781–12791 (2011).

24. A. M. Henao-Restrepo et al., Efficacy and effectiveness of an rVSV-vectored vaccine in preventing Ebola virus disease: final results from the Guinea ring vaccination, open-label, cluster-randomised trial (Ebola Ca Suffit!). Lancet 389, 505–518 (2017).

25. X. Qiu et al., Mucosal immunization of cynomolgus macaques with the VSVDeltaG/ZEBOVGP vaccine stimulates strong ebola GP-specific immune responses. PLoS One 4, e5547 (2009).

26. J. Emanuel et al., A VSV-based Zika virus vaccine protects mice from lethal challenge. Sci Rep 8, 11043 (2018).

27. V. J. Munster et al., Respiratory disease in rhesus macaques inoculated with SARS-CoV-2. Nature 585, 268–272 (2020).

28. A. Marzi et al., Antibodies are necessary for rVSV/ZEBOV-GP-mediated protection against lethal Ebola virus challenge in nonhuman primates. Proc Natl Acad Sci U S A 110, 1893–1898 (2013).

29. A. R. Menicucci et al., Transcriptome Analysis of Circulating Immune Cell Subsets Highlight the Role of Monocytes in Zaire Ebola Virus Makona Pathogenesis. Front Immunol 8, 1372 (2017).

30. J. Wang et al., Single mucosal, but not parenteral, immunization with recombinant adenoviral-based vaccine provides potent protection from pulmonary tuberculosis. J Immunol 173, 6357–6365 (2004).

31. M. R. Neutra, P. A. Kozlowski, Mucosal vaccines: the promise and the challenge. Nat Rev Immunol 6, 148–158 (2006).

32. L. H. A. Cavalcante-Silva et al., Neutrophils and COVID-19: The road so far. Int Immunopharmacol 90, 107233 (2020).

33. V. Francois-Newton et al., USP18-based negative feedback control is induced by type I and type III interferons and specifically inactivates interferon alpha response. PLoS One 6, e22200 (2011).

34. T. Matsumura et al., TIFAB inhibits TIFA, TRAF-interacting protein with a forkhead-associated domain. Biochem Biophys Res Commun 317, 230–234 (2004).

35. J. J. Wong, Y. F. Pung, N. S. Sze, K. C. Chin, HERC5 is an IFN-induced HECT-type E3 protein ligase that mediates type I IFN-induced ISGylation of protein targets. Proc Natl Acad Sci U S A 103, 10735–10740 (2006).

36. Y. Peng et al., Broad and strong memory CD4(+) and CD8(+) T cells induced by SARS-CoV-2 in UK convalescent individuals following COVID-19. Nat Immunol 21, 1336–1345 (2020).

37. O. Boyman, J. Sprent, The role of interleukin-2 during homeostasis and activation of the immune system. Nat Rev Immunol 12, 180–190 (2012).

38. G. Abrahamsen, V. Sundvold-Gjerstad, M. Habtamu, B. Bogen, A. Spurkland, Polarity of CD4+ T cells towards the antigen presenting cell is regulated by the Lck adapter TSAd. Sci Rep 8, 13319 (2018).

39. J. S. Woo et al., CRACR2A-Mediated TCR Signaling Promotes Local Effector Th1 and Th17 Responses. J Immunol 201, 1174–1185 (2018).

40. Y. Xiong et al., Transcriptomic characteristics of bronchoalveolar lavage fluid and peripheral blood mononuclear cells in COVID-19 patients. Emerg Microbes Infect 9, 761–770 (2020).

41. J. B. Case et al., Replication-Competent Vesicular Stomatitis Virus Vaccine Vector Protects against SARS-CoV-2-Mediated Pathogenesis in Mice. Cell Host Microbe 28, 465–474 e464 (2020).

42. A. Marzi et al., Single low-dose VSV-EBOV vaccination protects cynomolgus macaques from lethal Ebola challenge. EBioMedicine 49, 223–231 (2019).

43. A. Huttner et al., Determinants of antibody persistence across doses and continents after single-dose rVSV-ZEBOV vaccination for Ebola virus disease: an observational cohort study. Lancet Infect Dis 18, 738–748 (2018).

44. J. Harcourt et al., Severe Acute Respiratory Syndrome Coronavirus 2 from Patient with Coronavirus Disease, United States. Emerg Infect Dis 26, 1266–1273 (2020).

45. Y. Tsuda et al., Protective efficacy of a bivalent recombinant vesicular stomatitis virus vaccine in the Syrian hamster model of lethal Ebola virus infection. J Infect Dis 204 Suppl 3, S1090–1097 (2011).

46. H. B. Tw, T. Girke, systemPipeR: NGS workflow and report generation environment. BMC Bioinformatics 17, 388 (2016).

